# Longitudinal dynamics of organ-specific proteomic aging clocks over a decade of midlife

**DOI:** 10.64898/2026.02.17.706320

**Authors:** Robbe E. Neirynck, Julio A. Chirinos, Menno Van Damme, Louis Coussement, Patrick Segers, Marc L. De Buyzere, Ernst R. Rietzschel, Tim De Meyer

## Abstract

Organ-specific proteomic clocks are promising tools for quantifying heterogeneity in biological aging, but their longitudinal behavior remains largely unexplored. Here, we analyzed paired plasma proteomic profiles with 10-year follow-up in middle-aged adults (*n*= 1,250) to evaluate their longitudinal properties. Cross-sectional associations of protein concentrations with age mirrored average longitudinal trajectories, validating the common cross-sectional training of clocks. Organ-specific age acceleration was moderately stable over the decade, and aging across organs progressed in parallel, with the immune and adipose systems acting as central hubs and early cardiorespiratory aging predicting downstream metabolic aging. Critically, longitudinal changes in predicted age tracked subclinical risk factor alterations. In women, the menopausal transition dominated the aging landscape and was associated with multi-organ age acceleration. Medication initiation altered clocks through specific drug-targeted proteins (such as renin and APOB) rather than generalized organ aging. Together, these findings position organ-specific proteomic clocks as interpretable, dynamic indicators of aging and organ health.

## Introduction

Aging is characterized by a gradual loss of physiological integrity that drives functional decline and disease susceptibility^1^. The pace of this decline varies between individuals^2^ and even between organs within the same person^3,4^. This aging heterogeneity has led to the development of machine learning models that quantify “biological age”. Here, age is typically predicted from molecular profiles, and the difference between predicted age and chronological age (“age gap”) is used as a measure of biological aging. Over the past decade, a large range of omics assays have been used to construct such aging clocks^5–8^. Among these, proteomic clocks seem particularly compelling because proteins are the direct effectors of cellular function and offer greater biological interpretability, sitting at the center of many mechanisms that underlie cellular aging and disease. Moreover, as many proteins are preferentially produced by specific tissues, their blood plasma concentrations can be used as a minimally invasive strategy to build organ-specific clocks^4^. This approach is supported by recent studies showing that accelerated organ aging predicts future organ-specific morbidity better than conventional aging clocks^4,9,10^. Consequently, these clocks have been proposed as tools for monitoring organ health and for evaluating anti-aging interventions.

Despite their promise, organ-specific proteomic clocks have been evaluated almost exclusively in cross-sectional, single-measurement settings. Even at the level of individual proteins, large-scale longitudinal analyses are scarce or involve only a limited subset of proteins^11^. As a result, it is unclear whether changes in organ clocks track meaningful changes in organ health over time or whether they function mainly as static baseline predictors. This distinction matters, because dynamic and potentially modifiable behavior would extend their utility from risk stratification to active monitoring of physiological organ health. Moreover, only multi-timepoint data can show how organ age gaps evolve within individuals and how aging patterns propagate across organs.

To address these questions, we conducted a longitudinal analysis of 6,402 circulating plasma proteins measured at two time points approximately a decade apart in 1,250 generally healthy middle-aged participants. We first characterized age trajectories of individual proteins and found that cross-sectional associations serve as a good proxy for within-individual changes. We then developed and evaluated both general and sex-specific proteomic clocks for eleven organs following the framework of Oh et. al^4^. We show that organ age gaps are relatively stable within individuals and that acceleration in one organ often coincides with acceleration in others. The paired measurements further enabled us to study the temporal propagation of biological aging across organs. Finally, we tested whether changes in organ age gaps track concurrent changes in clinical health markers, lifestyle factors, menopausal transition, and medication initiation, thereby assessing the value of these gaps as dynamic health monitoring tools.

## Results

### Cross-sectional protein aging mirrors within-person trajectories

The cohort consisted of 1,250 adults free of overt cardiovascular disease at baseline. Phenotype summary statistics are provided in Extended Data Table 1. After quality control and data preprocessing (Methods), 1,128 baseline samples (mean age 46 years) and 1,228 follow-up samples (mean age 56 years) were retained. In total, 1,114 participants (50.9% female) had successful plasma protein measurements at both visits, with a mean interval of 9.9 ± 1.5 years between visits (Fig. 1a; Extended Data Fig. 1).

**Fig. 1:**
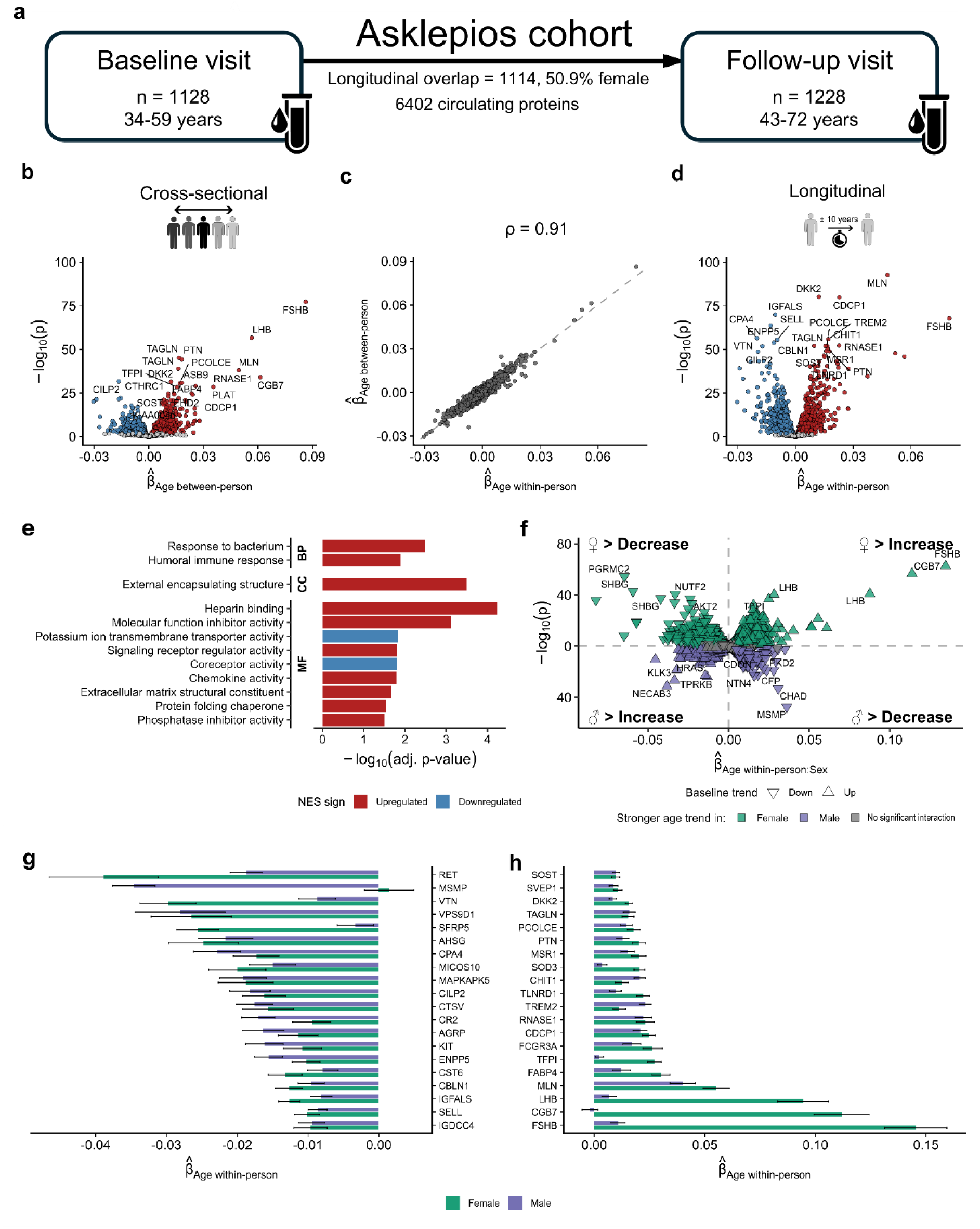
Cross-sectional protein–age associations reflect true aging changes and differ between sexes. **a**, Overview of the longitudinal protein measurements in the Asklepios cohort. After quality control, 1128 baseline and 1228 follow-up samples remained, resulting in 1114 participants with proteomic data from both visits. **b–d**, Longitudinal changes in circulating plasma proteins (7,289 aptamers) were analyzed in 1,114 individuals with measurements at both visits using a within–between mixed-effects framework that included sex as a covariate and a random intercept for each individual. **b**, Volcano plot of between-person aging effects. **c**, Spearman correlation between within-person and between-person effects. **d**, Volcano plot of within-person aging effects. **e**, Gene set enrichment analysis of proteins ranked by significance of the within-person aging effects, using Gene Ontology (GO) and KEGG pathways (no KEGG pathways were significantly enriched). NES = normalized enrichment score. **f**, Interaction between the within-person aging effect and sex. Points above the y=0 line indicate steeper age-related trends in females, and points below the line steeper trends in males. Point shape denotes the sign of the baseline within-person aging effect in the model without the interaction. **g, h**, Sex-specific within-person aging effects for the top 20 decreasing (f) and increasing (g) proteins, obtained by fitting the model separately in men and women. Error bars represent 95% confidence intervals.

Most proteomic aging studies, including those developing clocks, rely on cross-sectional data and implicitly assume that between-person differences by age directly reflect within-person aging. Therefore, we first tested whether these cross-sectional age associations mirror true longitudinal change. Of 6,402 proteins assessed, 2,446 showed significant within-person age associations over the decade (52.5% decreasing, 47.7% increasing) and 1,266 showed significant between-person associations with age (43.0% decreasing, 57.0% increasing) (Supplementary Table 1). Importantly, effect estimates were highly correlated between analyses (ρ = 0.91; Fig. 1b–d). Thus, although the longitudinal analysis provided greater power to detect age trends, cross-sectional coefficients were a good proxy for within-person proteomic change over midlife, with limited evidence for generational bias.

In within-person analyses, proteins increasing with age were functionally most enriched for heparin binding (Fig. 1e), consistent with prior cross-sectional proteomic reports implicating coagulation and other blood-related pathways in aging^5,12^. Conversely, proteins decreasing with age were most enriched for potassium ion transmembrane transporter activity (Fig. 1e), possibly reflecting known age-related reductions in potassium channel expression and Na⁺/K⁺-ATPase activity^13,14^.

### Half of age-related proteins show sex-dependent trajectories

In this middle-aged cohort, nearly half of longitudinally age-associated proteins featured a significant sex interaction (1,122; 45.9%; Fig. 1f; Supplementary Table 2 and 3). This finding aligns with both proteomic and broader studies suggesting that males and females age differently^12,15^. For most of these proteins (87.2%), the magnitude of the age effects was larger in females. This female dominance is not universal across cohorts and likely depends on the age window examined. For instance, a study in an older cohort (age range: 65-95 years) reported a greater number of proteins changing with age in men^16^. In Asklepios, follow-up spans on average from 45 to 55 years, which overlaps the typical menopausal transition and likely explains the larger age-related changes observed in women. Accordingly, two of the strongest age-increasing proteins were the menopause-related gonadotropins follicle-stimulating hormone (FSHB) and luteinizing hormone (LHB) (Fig. 1h). Of note, we observed a pronounced age-related decline in prostate-associated microseminoprotein (MSMP) in men but not in women (Fig. 1g). Surprisingly, we could not find prior reports describing age-related change in circulating MSMP, even though MSMP has recently been identified as one of the top protein markers associated with lower frailty risk^17^. Altogether, these widespread sex-dependent age trajectories motivated us to train both sex-combined and sex-stratified clocks.

### Clock performance is on par with the state of the art

Using 16,245 bulk RNA-seq samples from 51 human tissues^18^, we defined mutually exclusive sets of organ-enriched proteins as those with at least four-fold higher expression in one organ system compared with all others^4^ (Methods and Supplementary Table 4). We then trained proteomic aging clocks for 11 organ systems using SomaScan 7K samples from both visits (n = 2,356), following the framework to Oh et al^4^ with minor modifications (Methods).

Across the general (sex-combined) models, out-of-fold performance ranged from r = 0.91 for the full-body clock to r = 0.31 for the lung clock (MAE = 2.55–6.06 years; Fig. 2a), on par with the performance reported by Oh et al.^4^. When applied to the Asklepios cohort, the Oh et al. released SomaScan 5K clocks had a lower accuracy than our models for all organs, except for the intestines (Fig. 2a).

**Fig. 2:**
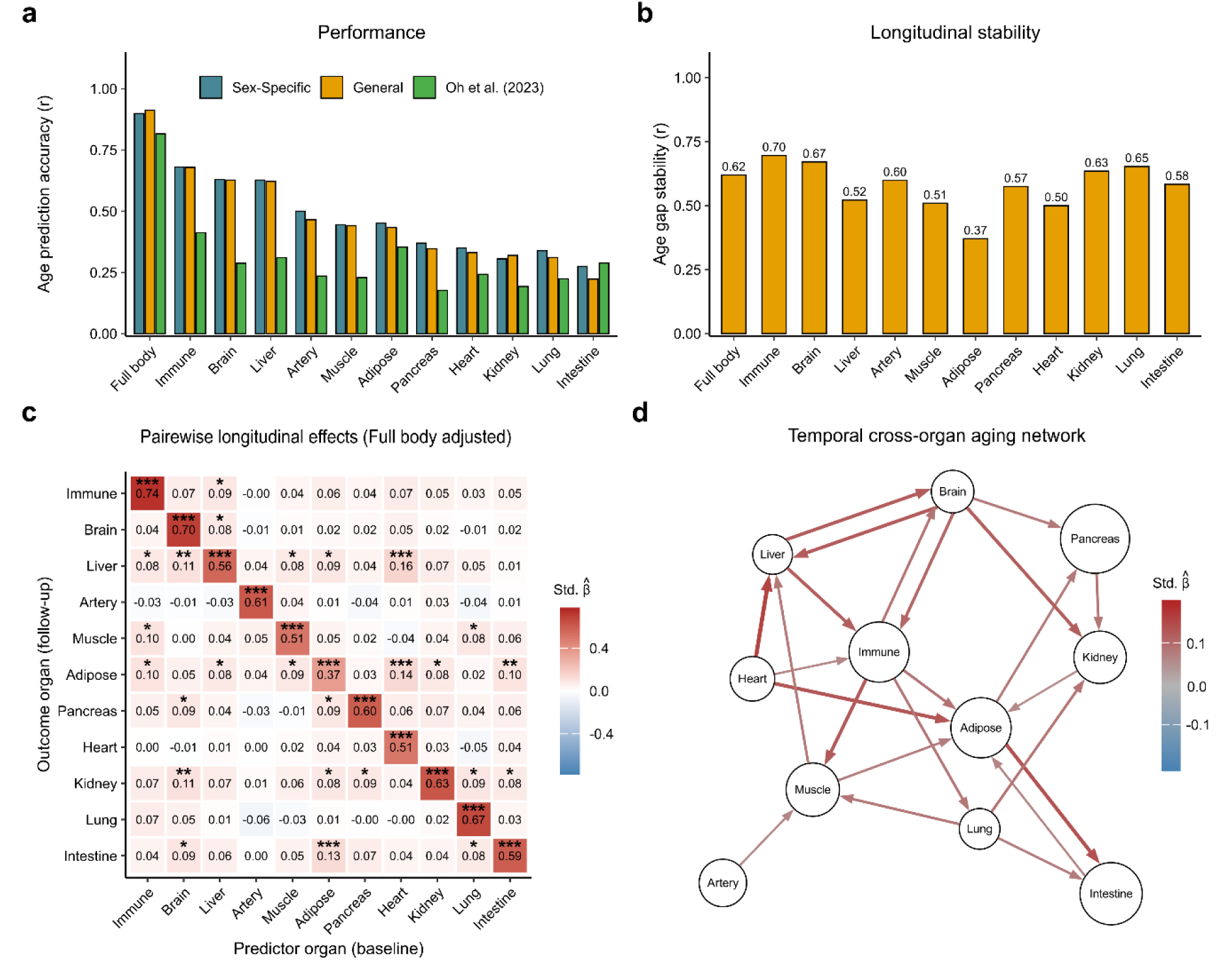
Longitudinal dynamics and cross-organ aging effects. **a**, Comparison of out-of-fold age-prediction performance across organs for thQree models: a sex-specific clock, a general clock, and the Oh et al. (2023) clock. **b**, Longitudinal stability of the general organ clock, calculated as the correlation of z-scored age gaps between baseline and follow-up for each organ. **c**, Pairwise cross-organ longitudinal associations corrected for systemic aging. Follow-up age gaps were predicted from baseline age gaps (z-scored), adjusted for sex and baseline full-body age gap (z-scored). P values were FDR-adjusted (*Adj. p < 0.05, **Adj. p < 0.01, *Adj. p < 0.001). **d**, Consensus cross-organ aging network obtained using a causal-discovery approach applied to sex-adjusted baseline and follow-up age gaps. Edges were restricted to baseline-to-follow-up directions and self-edges were excluded.

Sex-stratified training only marginally increased accuracy, with small improvements in 10 of 11 organ models and a slight decrease for the full-body clock (Fig. 2a). When evaluated within each sex, general and sex-stratified models performed surprisingly similar across organs (Extended Data Fig. 2). Also, both consistently achieved higher accuracy in females, in line with our finding above that most sex-interacting proteins had larger, and thus more informative, age effects in women. The important coefficients from the male- and female-stratified clocks were only moderately correlated (mean ρ = 0.55, range 0.36–0.88 across organs; Extended Data Fig. 3; summaries in Supplementary Table 6 and 7), pointing to a shared age signal but also clear sex differences. For instance, the female full-body clock showed significantly enriched contributions from endocrine proteins (FSHB, LHB, PRL, INHBA/INHBC, PPY and NPPB) (Extended Data Fig. 4). Nevertheless, given the comparable performance and the prior clinical validation of general organ clocks, we focused subsequent longitudinal analyses on the dynamics of the latter models.

### Organ aging differences persist over time and predict future aging of other organs

The age-adjusted difference between predicted and chronological age (the “age gap”) is used to quantify whether an organ appears older or younger than same-aged peers. Baseline organ age gaps have been linked to organ health and future disease risk^4,9^, with a recent validation study showing that immune, liver, heart, and lung clocks selectively predict disease risk in their corresponding organs^10^. Their within-individual evolution, however, has not been comprehensively examined yet.

In this cohort, organ age acceleration persisted over the 10-year follow-up. Organs that appeared older at baseline tended to remain relatively older at follow-up, with a median Pearson r = 0.58 between baseline and follow-up age gaps across organ clocks (range 0.37–0.70; Fig. 2b). We also observed that longitudinal changes in organ age gaps were broadly coordinated: changes in the age gaps were significantly positively correlated in 54 of 55 pairwise organ combinations after multiple-testing correction (median pairwise Pearson r = 0.21). This shows that within-individual acceleration in one organ typically co-occurs with acceleration in others, reflecting a shared systemic aging component. Adjusting these analyses for the baseline full-body age gap indeed attenuated most cross-organ associations, yet a subset of baseline-to-follow-up cross-organ links persisted (26/110 possible directed organ pairs; Fig. 2c).

To explore whether baseline biological age in one organ influenced future biological age in others, we applied a temporally constrained causal discovery framework^19^ and derived a consensus network of directed cross-organ links (Fig. 2d; Methods). All final cross-organ edges were positive, reflecting reinforcing aging dynamics across organ systems. Three organ pairs had bidirectional links—liver–brain, brain–immune, and adipose–intestine—forming direct feedback loops within these subsystems. Adipose and immune occupied central strongly connected positions in the network and, together with muscle, resembled an immunometabolic “fitness” axis. The heart emerged as a prominent upstream node, with baseline heart age gap predicting higher biological age at follow-up in the liver and adipose tissue. Although the heart-to-adipose directionality may appear counterintuitive, but could be plausibly explained by the endocrine role of B-type natriuretic peptide (BNP/NPPB; the largest contributor to the heart clock; Extended Data Fig. 4), which promotes adipose lipolysis and thermogenic browning^20,21^. Lung aging likewise showed predominantly outgoing links to kidney, muscle, and intestine, while being influenced by early immune aging.

### Changes in organ age gaps track clinical profiles and the menopausal transition

We next examined how organ age gaps relate to clinical risk factors and subclinical phenotypes. Consistent with prior reports of faster biological aging in men^9,22^, age gaps were on average higher in males for 8 of 11 organs (Extended Data Fig. 5). Within each sex, however, phenotype associations derived from general and sex-specific clocks were highly similar (Extended Data Fig. 6 and 7), showing that sex-specific training does not materially change clinical interpretation.

Cross-sectionally, cardiometabolic and renal markers such as higher BMI, glucose, homocysteine, fibrinogen, and uric acid, as well as reduced kidney function (lower eGFR and higher creatinine), were associated with accelerated aging across most organs (Fig. 3a). Importantly, many of these associations also held within individuals. Over ten years, increases in these clinical markers tended to track with worsening organ age gaps (Fig. 3b). For example, individuals who developed diabetes showed accelerated aging of the kidney, pancreas, and intestine, while increases in biological heart age were related to higher left-ventricular mass, higher stroke volume, and lower resting heart rate. Overall, these findings support the potential of organ clocks as dynamic indicators of organ health.

**Fig 3.**
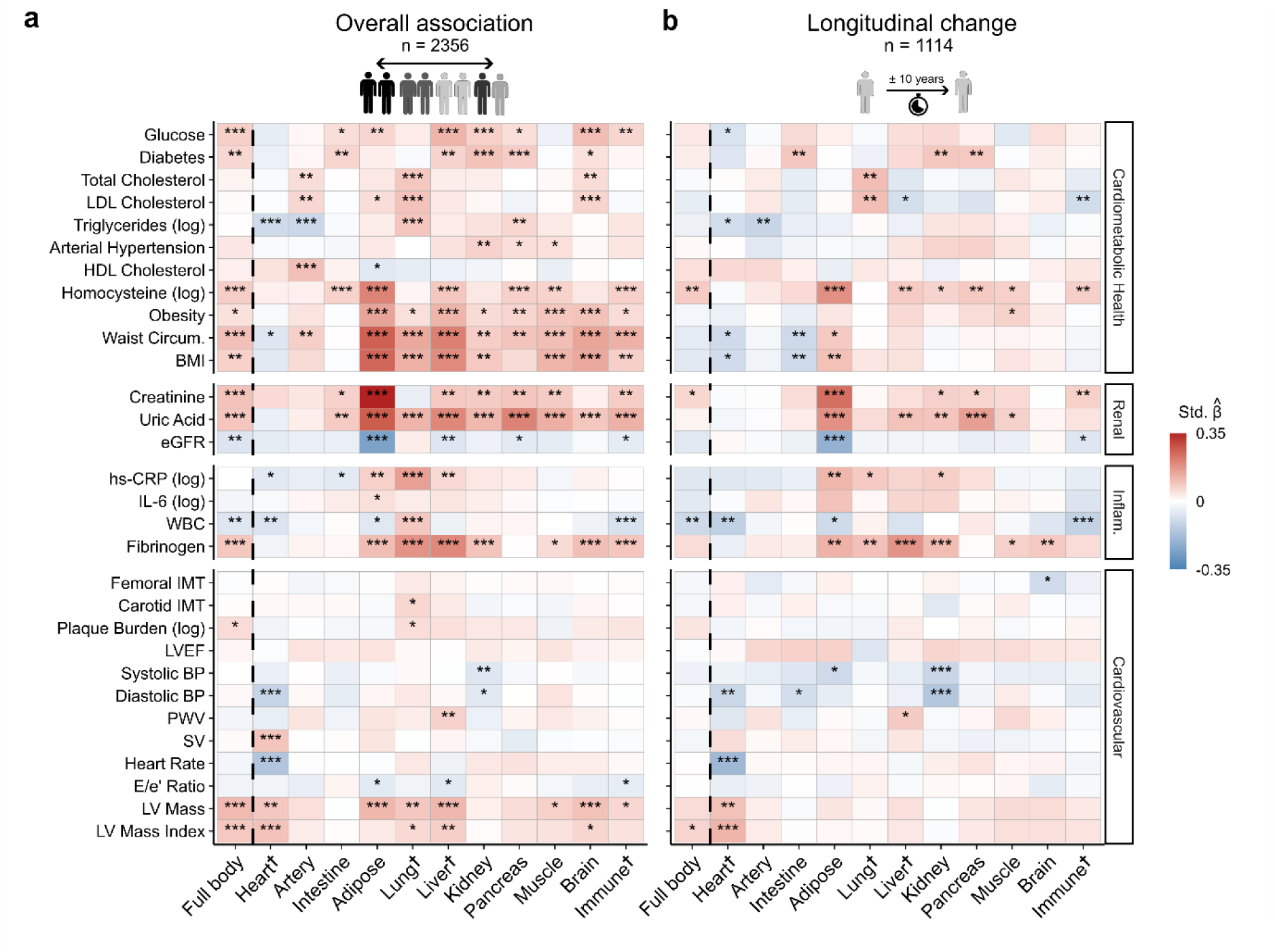
Organ age gaps track clinical profiles and their 10-year change. **a**, Cross-sectional associations between general organ age gaps and cardiometabolic, renal, inflammatory, and cardiovascular markers across all samples (n = 2,356; 1,128 baseline and 1,228 follow-up). Linear mixed-effects models regressed z-scored organ age gaps on z-scored clinical markers, adjusted for age and sex with a random intercept for participant. **b**, Longitudinal associations between (approximately) 10-year changes in organ age gaps and changes in the same markers (n = 1,114 individuals with proteomic data at both visits). Linear models predicted follow-up z-scored organ age gaps from z-scored biomarker changes, adjusted for z-scored baseline age gap, z-scored baseline biomarker level, mean age, time between visits, and sex. P values were FDR-adjusted separately for the cross-sectional and longitudinal analyses (*Adj. p < 0.05, **Adj. p < 0.01, ***Adj. p < 0.001). **†**Organs with clocks that have strong independent validation for uniquely predicting diseases specific to their respective organ^10^. Abbreviations: BMI, Body Mass Index; BP, Blood Pressure; eGFR, estimated Glomerular Filtration Rate; hs-CRP, high-sensitivity C-reactive protein; IL-6, Interleukin-6; IMT, Intima-Media Thickness; LV, Left Ventricular; LVEF, Left Ventricular Ejection Fraction; NSAID, Non-Steroidal Anti-Inflammatory Drug; PWV, Pulse Wave Velocity; RAS, Renin–Angiotensin System; SV, Stroke Volume; WBC, White Blood Cell count.

Notably, decreases in total white blood cell (WBC) count were associated with an increase in biological immune age over the decade, a pattern also seen for the full-body clock. This relationship was almost entirely driven by women, whose WBC counts declined more steeply with age (r = –0.25) than those of men (r = –0.05) at follow-up (Extended Data Fig. 8). Among the WBC subfractions in women, neutrophils showed the strongest age-related decline (r = –0.23), but rising FSH levels were an even stronger correlate of lower neutrophil counts (r = –0.36) than age itself (Extended Data Fig. 8). Together, this suggests that the WBC–immune-age association in women primarily reflects the known menopause-related reduction in neutrophils^23^. Guided by this observation, we investigated the dynamics of the menopause status, which we defined by clustering women based on their FSH levels (Fig. 4a-b). Here we found that women who transitioned from pre-to post-menopause during follow-up (n = 231) indeed showed significantly accelerated immune aging compared with stably pre-menopausal (n = 234) or stably post-menopausal (n = 118) groups (Fig. 4c-f). Similar effects in the full-body, liver, and artery clocks point to broader biological age acceleration following the menopausal transition.

**Fig. 4:**
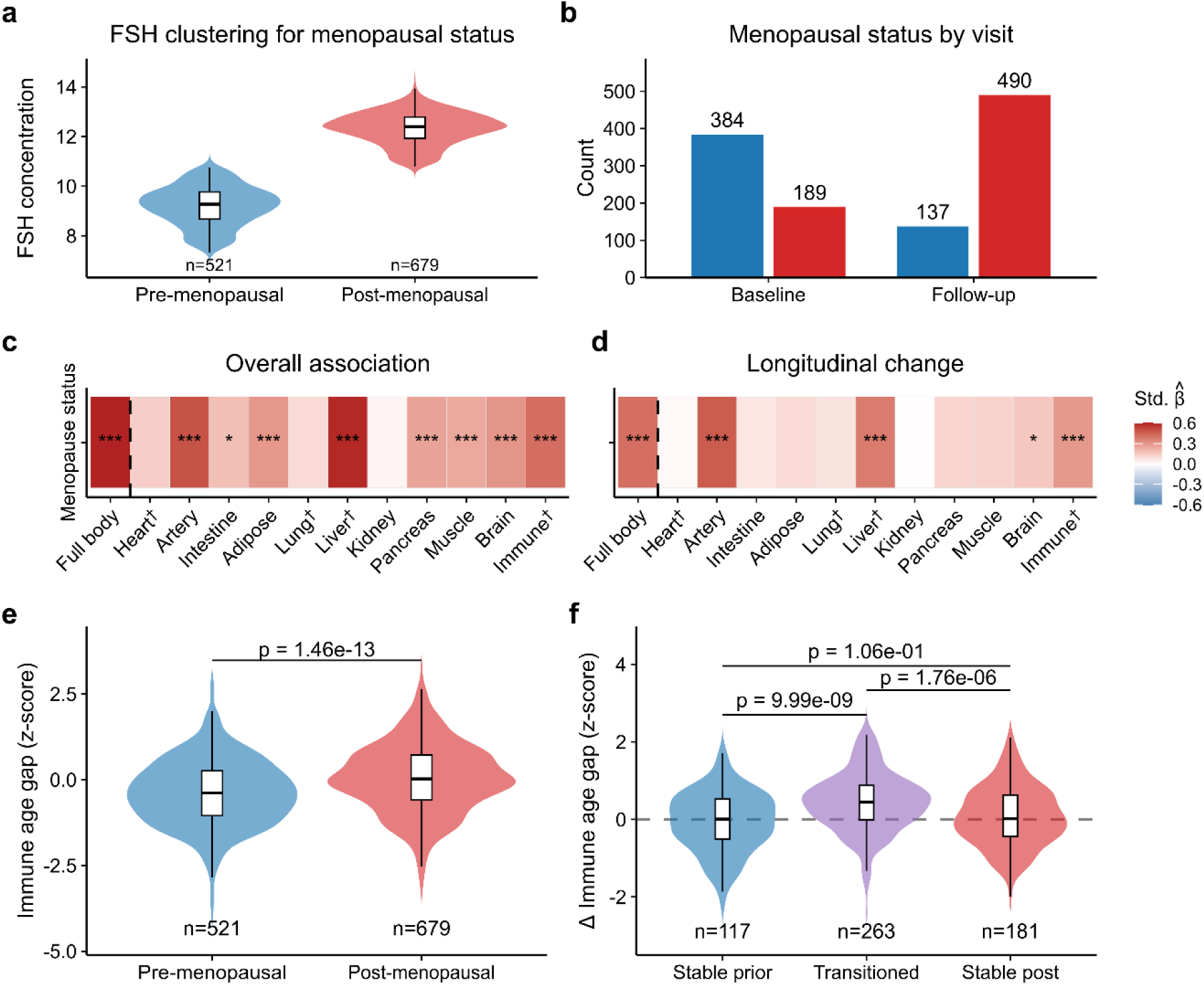
Menopausal transition accelerates biological aging across multiple organs. **a, b**, Determination of menopausal status in female participants (n = 1,200 samples) using K-means clustering (k = 2) of plasma follicle-stimulating hormone (FSH) concentrations. **a**, Distribution of FSH concentrations by cluster-defined status. **b**, Participant counts in the prior- and post-menopause groups at baseline and follow-up visit. **c**, Cross-sectional associations between menopausal status (post-versus prior-menopause) and z-scored organ age gaps. Coefficients are derived from linear mixed-effects models adjusted for chronological age with a random intercept for participant. (*Adj. p < 0.05, **Adj. p < 0.01, ***Adj. p < 0.001). **d**, Longitudinal associations between the menopausal transition and changes in organ age gaps in the subset of women with paired data (n = 561). Linear models predicted the follow-up organ age gap (z-scored) from the change in menopausal status (transitioned vs. stable), adjusted for the baseline age gap (z-scored), mean age, and time between visits. **e**, Distribution of immune age gaps (z-scored) by menopausal status (combined visits). P-values are derived from unpaired two-sided t-tests. **f**, The approximately 10-year change in immune age gap (Δ z-score) stratified by transition trajectory. Women who transitioned into the cluster-defined post-menopausal state (n = 231) experienced accelerated aging compared to those who remained stably prior-menopausal (n = 234) or stably post-menopausal (n = 118). †Organs with clocks that have strong independent validation for uniquely predicting diseases specific to their respective organ^10^.

### Medication trajectories, but not lifestyle changes, found to influence organ age gaps

Beyond tracking clinical markers, an important open question concerning organ clocks is whether the age gaps are modifiable by changing lifestyle or medication usage. As reported previously^9,24^, we also observed strong cross-sectional associations between lifestyle factors and organ age gaps, such as higher lung age in smokers and higher pancreas age in heavy drinkers (Fig. 5a). However, longitudinal changes in smoking or alcohol consumption were not linked to changes in organ age gaps (Fig. 5b), possibly reflecting limited natural lifestyle alterations in this general cohort.

**Fig. 5:**
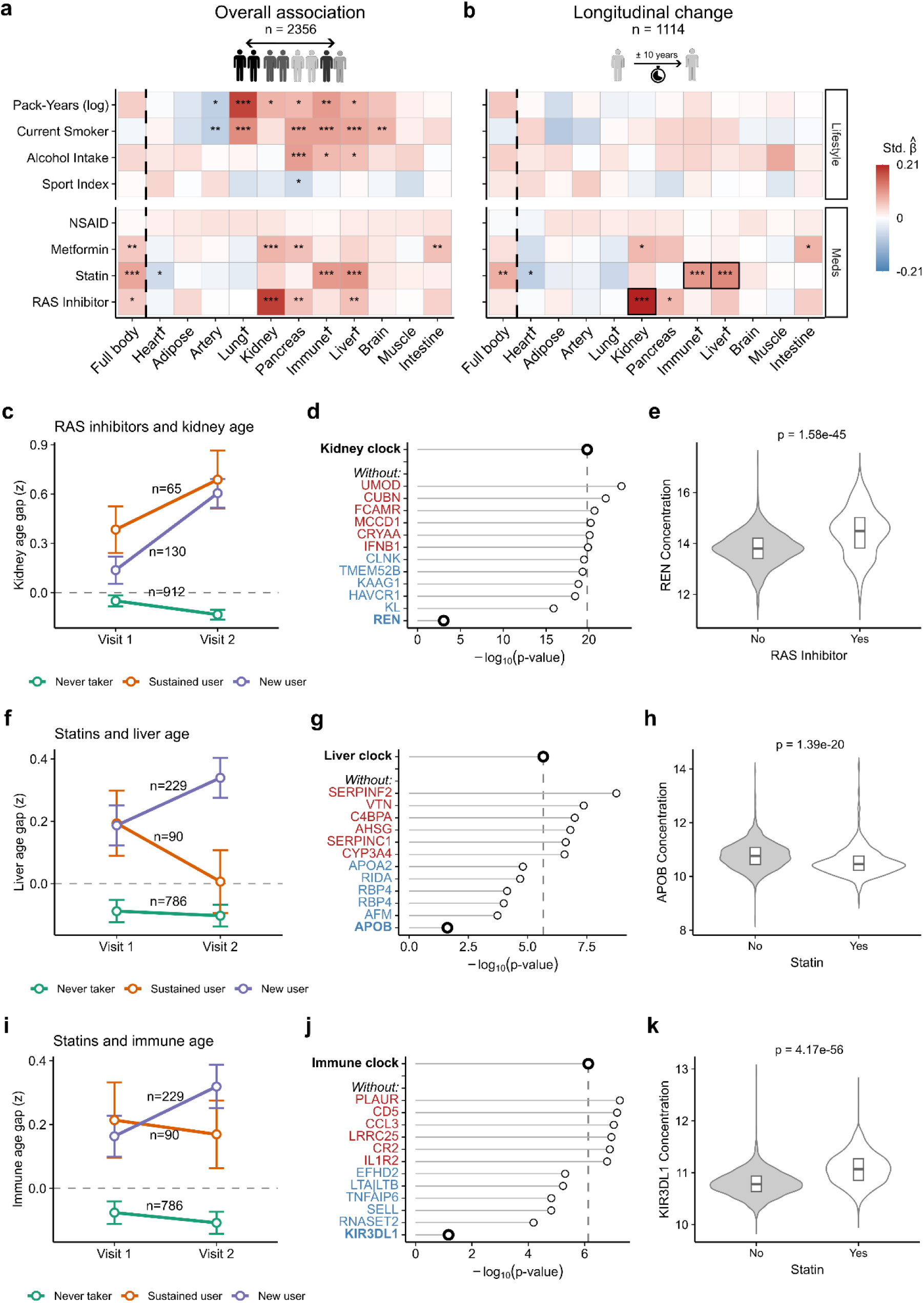
Medication trajectories shape organ age gaps through individual protein drivers. **a**, Cross-sectional associations between general organ age gaps and lifestyle and medication data across all samples (n = 2,356; 1,128 baseline and 1,228 follow-up). Linear mixed-effects models regressed z-scored organ age gaps on z-scored phenotypes, adjusted for age and sex, with a random intercept for participant. **b**, Longitudinal associations between (approximately) 10-year changes in organ age gaps and changes in the same phenotypes (n = 1,114 individuals with proteomic data at both visits). Linear models predicted follow-up z-scored organ age gaps from z-scored phenotype changes, adjusted for z-scored baseline age gap, z-scored baseline phenotype level, mean age, time between visits, and sex. P values were FDR-adjusted separately for the cross-sectional and longitudinal analyses (*Adj. p < 0.05, **Adj. p < 0.01, ***Adj. p < 0.001). **c, f, i**, Longitudinal trajectories of z-scored organ age gaps from baseline to follow-up. Kidney age gaps are shown by RAS inhibitor trajectories and liver and immune age gaps by statin trajectories. “New user” denotes participants who initiated medication between visits. Points represent group means ± standard error. Discontinuers of RAS inhibitors (n = 7) and statins (n = 9) were excluded from the visualization owing to small sample size. **d, g, j**, Top six positive and negative individual proteins driving the longitudinal kidney, liver, and immune age-gap associations with RAS inhibitor and statin use, respectively. The x-axis represents the significance of the association between z-scored change in medication use and follow-up z-scored organ age gap, from linear models adjusted for z-scored baseline age gap, z-scored baseline medication status, mean age, time between visits, and sex. Each “without” point represents an organ aging clock in which the coefficient(s) of that specific protein were set to zero, the age gaps were recalculated and the longitudinal association model was retested. Proteins whose removal weakens the association (blue) are considered positive drivers, while those whose removal strengthens it (red) are negative drivers. **e, h, k**, Violin plots comparing the distributions of REN, APOB, and KIR3DL1 concentrations by RAS inhibitor status (No/Yes, n = 1,049/72 at Visit 1; 1,017/211 at Visit 2) and statin status (No/Yes, n = 1,022/99 at Visit 1; 864/364 at Visit 2), respectively. Analyses included all participants with successful protein measurements and no missing data at Visit 1 (n = 1,121) and Visit 2 (n = 1,228). Annotated p-values are from linear mixed-effects models adjusted for age, sex, BMI, and eGFR, with a random intercept for each participant.

In contrast, the initiation of common cardiometabolic medications did show significant associations. Statin initiation during follow-up was linked to rising immune and liver age gaps, and renin–angiotensin system (RAS) inhibitor initiation to increasing kidney age gaps (Fig. 5cfi). For RAS inhibitors, their use was strongly associated with higher renin concentrations (Fig. 5e), consistent with the known compensatory increase in renin under RAS blockade^25^. When we set the renin coefficient in the kidney clock to zero, the association between RAS-inhibitor initiation and the kidney age gap was strongly attenuated (p from 1.62 × 10⁻²⁰ to 8.9 × 10⁻⁴), but not abolished, suggesting a modest effect beyond renin itself (Fig. 5d). As a side note, the surprising longitudinal finding in Fig. 3b—that increasing blood pressure was associated with a decrease in kidney age gaps—was also entirely carried by renin, although it remained significant after adjustment for medication use.

For statins, we unexpectedly observed that initiation was related to higher liver and immune age gaps. Sensitivity analysis showed that the liver association was largely driven by apolipoprotein B (APOB) (Fig. 5f), the main structural protein of LDL particles lowered by statins, as the liver association became only marginally significant after setting the APOB coefficient to zero (p = 0.023) (Fig. 5g). For the immune clock, the association was mainly driven by KIR3DL1 (Fig. 4j), an inhibitory receptor with previously reported immune-modulatory responses to statin therapy^26^; after removing KIR3DL1, the statin–immune age association was no longer significant (p = 0.068). Together, these findings illustrate that apparent “accelerations” of organ age can be driven by drug-induced changes in single proteins, rather than a generalized worsening of organ health per se.

## Discussion

This study provides the first comprehensive longitudinal evaluation of organ-specific proteomic clocks, performed through the follow-up of 6,402 proteins over 10 years in 1,250 carefully characterized middle-aged participants from the general population. Recent work has shown that these clocks predict future organ disease and mortality beyond standard risk factors and clinical markers^4,9,10,24^, but has explicitly called for longitudinal data to test the stability and modifiability of organ-specific age gaps, to clarify the specific order of organ aging, and to distinguish baseline differences from the dynamic tracking of phenotypes.

Here, we show that organ age gaps are moderately stable over ten years, with baseline values explaining on average one third of the variance at follow-up. Individuals who aged faster in one organ also tended to age faster in others, consistent with a shared systemic aging component. Yet, our causal discovery analysis further supported a temporal hierarchy in cross-organ aging. At the network’s center, adipose and immune aging connected to most other systems, in line with the central role of chronic low-grade inflammation in multisystem aging^27^ (inflammaging) and with the view of adipose tissue as an active endocrine organ that shapes whole-body metabolic state rather than a passive energy reservoir^28^. The heart and lung emerged as predominantly upstream nodes, where early cardiorespiratory aging was linked to subsequent aging in metabolic organs including the liver, intestine, kidney, and adipose tissue, supporting the idea that improving cardiovascular and respiratory function could yield downstream benefits for slowing proteomic aging in dependent organs.

Importantly, we found that acceleration of organ age over the decade tracked with worsening cardiometabolic and renal markers (e.g., fibrinogen, homocysteine, uric acid, and creatinine). For instance, individuals who developed type 2 diabetes showed an increase in the biological ages of the pancreas, kidney, and intestine—organs intimately involved in metabolic regulation. These results suggest that proteomic clocks capture not just static risk, but the active trajectory of aging. This dynamic sensitivity was perhaps most visible during the menopausal transition, one of the strongest modifiers of biological aging in our study. Women who moved from pre-to post-menopause during follow-up showed clear acceleration of immune aging, alongside increases in full-body, liver, and arterial age gaps. These shifts paralleled declines in white blood cell counts, particularly neutrophils, a well-known consequence of falling estrogen levels^23^. Moreover, two of the most age-associated proteins in our dataset were markers of menopause (FSH and LH). This, alongside the enrichment of hormone-related proteins in the female full-body clock, show how the menopausal transition dominates the proteomic landscape in midlife women. Together, these findings indicate that loss of ovarian function accelerates molecular aging across multiple organs and suggest that targeting this transition, potentially through hormone replacement therapy^29^, may be important to slow down aging in midlife women.

More broadly, aging proved highly sex-dependent: nearly half of all age-related proteins showed significant sex interactions. Nevertheless, sex-specific clocks did not meaningfully outperform general models or alter their phenotypic associations. This suggests that while men and women may age through partially distinct molecular pathways, the end point—in terms of organ biological age relative to health—is comparable. Future clock designs may benefit from more complex architectures that allow for sex interactions, though such approaches will likely require larger datasets than were available here.

The importance of interpretability in biological clocks was clearly illustrated by our analysis of medication trajectories. Although initiation of statins and RAS inhibitors appeared to accelerate liver and kidney aging, these signals could be traced to single drug-targeted proteins (APOB and renin) that were also major contributors to their respective clocks. Removing these proteins from the models strongly attenuated the associations, indicating that the clocks were capturing the expected statin-induced LDL lowering and the compensatory rise in renin under RAS blockade^25^, rather than a generalized acceleration of organ aging. This level of transparency stands in sheer contrast to most methylation clocks, which rely on large sets of CpG sites chosen primarily for statistical performance. Different methylation clocks show only very limited overlap in CpG composition^30^, and the functional relevance of many loci remains unclear^31^. As a result, shifts in epigenetic age can be difficult to interpret mechanistically and may not always reflect true biological aging.

Finally, we recognize several limitations. First, the Asklepios cohort’s healthy middle-aged profile meant we had few hard clinical endpoints (e.g., disease onset) to relate to organ clock changes. In the context of preventive monitoring, however, it remains important to assess whether these clocks track subclinical variation and early deviations from healthy trajectories. On a related note, while we do see clear cross-sectional effects, the absence of significant change of lifestyle effects likely reflects the limited natural variation in lifestyle within a general middle-aged cohort rather than true non-modifiability through lifestyle. Controlled intervention studies, like those demonstrating reversal of epigenetic age with diet and lifestyle changes^32^, will be needed to fully prove modifiability through lifestyle alterations. Second, the two-timepoint design, although a major advance over cross-sectional data, cannot capture short-term fluctuations or the non-linear aging trajectories that characterize many proteins across the lifespan^12^. Third, our organ clocks were trained to predict chronological age rather than morbidity or functional decline. While this follows the paradigm of most organ-specific proteomic clock studies, maximizing age prediction does not necessarily yield the most clinically informative biomarker^33^. Moreover, because proteomic signatures of healthy aging can diverge from those of age-related disease (e.g., SCGB1A1 increases with normal lung aging but decreases in COPD^9,34^), future generations of clocks trained on clinical endpoints or functional decline may prove more translatable. Fourth, although the temporal ordering in the cross-organ network precludes reverse causation, confounding and the small magnitude of cross-organ effects relative to within-organ stability warrant an exploratory interpretation of the network. Finally, although we addressed batch and other technical biases as rigorously as possible (Supplementary Methods Note), SOMAmer measurements can still be influenced by off-target binding or by genetic variants that alter aptamer binding independent of true protein concentration.

In summary, this study provides the first longitudinal validation of organ-specific proteomic clocks. Beyond confirming their stability over time, we show how organ aging cascades through a temporal hierarchy. The ability of these clocks to dynamically track subclinical risk factors and the menopausal transition highlights their potential to detect early deviations from healthy aging trajectories. Our medication analyses further illustrate that clinically useful clocks must be interpretable and ideally decomposable. Taken together, these models represent a powerful, minimally invasive toolkit for quantifying an actionable biological organ age.

## Methods

### Asklepios Cohort

The rationale, design and exclusion criteria of the Asklepios study have been described previously^35^. Briefly, at baseline, apparently healthy individuals without overt cardiovascular disease were randomly recruited from the Belgian twin communities of Erpe-Mere and Nieuwerkerken. The Ghent University hospital ethics committee approved the study protocol, and participants provided written informed consent (baseline approval number 2002/133; follow-up approval B670201111695).

### Proteomic profiling

From the 2,254 Asklepios participants, a random subset of 1,250 individuals with blood plasma samples at both baseline and follow-up was selected for plasma proteomic profiling using the SomaScan 7K assay. Baseline samples were measured in 2022 and follow-up samples in 2023. Laboratory procedures for proteomic profiling have been described in detail elsewhere^36^. Of the 7,523 slow off-rate modified aptamers (SOMAmer reagents) on the SomaScan 7K platform, we retained the 7,289 reagents targeting human proteins. We excluded samples based on SomaLogic quality control flags (n = 96 baseline, n = 17 follow-up), as well as deviating samples identified by outlying age (n = 3 individuals, excluded at both timepoints), aberrant intensity distributions (n = 1 baseline, n = 1 follow-up), and outlier status in UMAP analysis (n = 22 baseline). After quality control, 1,128 baseline and 1,228 follow-up samples remained, with 1,114 participants having paired measurements. We further preprocessed the proteomic data to address technical variation related to batch differences between both visits, sample storage time, and plate effects; full details on quality control and preprocessing are provided in the Supplementary Methods Note.

### Protein-level age trajectories

To model how individual proteins change with age, we fitted a within–between mixed-effects model for each of the 7,289 aptamers using data from the 1,114 participants with repeated measures:

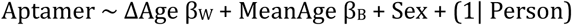

where MeanAge is the average age of a participant across both visits, ΔAge = Age – MeanAge, and Person is a random intercept for each participant. This model allows for unbiased disaggregation of age variance into longitudinal (within-person, β_W_) and cross-sectional (between-person, β_B_) associations^37^. To evaluate sex interactions on a longitudinal scale, we added a ΔAge × Sex term. Models were fitted using the lmerTest R package^38^. In the Results, a protein (identified by its UniProt ID) was reported as significantly associated with age if at least one of its corresponding aptamers passed the global Benjamini–Hochberg FDR threshold of 5%; for these proteins, directionality was determined by the aptamer with the most significant age relationship.

### Gene set enrichment analysis

We performed gene set enrichment analysis (GSEA) to identify pathways enriched for longitudinal age-related protein changes. Aptamers were mapped to Entrez gene symbols; for multi-gene annotations (e.g., “CGA|FSHB”), only the final, more specific symbol was used. Genes were ranked by the signed t-statistic of their within-person age coefficient. When multiple aptamers mapped to the same gene, the aptamer with the largest absolute t-statistic was used. This ranked list was analyzed with the fgsea R package^39^ against Gene Ontology (GO) and KEGG collections from MSigDB^40^, restricting gene sets to a size of 15–500 genes. Redundant pathways were collapsed using the collapsePathways function in fgsea^39^.

### Identifying organ-specific proteins

Raw gene-level read counts from GTEx v8 were available for 51 tissue regions (totaling 16,245 samples)^18^. Genes with missing values in any sample were removed. Counts were normalized using the trimmed mean of M-values (TMM) method in edgeR^41^, and converted to counts per million (CPM). We retained only genes mapping to SomaScan 7K aptamers via Ensembl IDs (n = 6,319). Tissue-level median CPM was computed per gene and aggregated to 21 organ systems following Oh et al.^4^ by taking the maximum median expression among tissues within each organ. Genes were classified as organ-enriched if expression in the top organ was more than fourfold higher than in the second-highest organ. To exclude proteins with high technical variability, we removed those whose assay coefficient of variation (CV) exceeded 13.5%, based on SomaScan 7K variability assessments^42^. The “Full body” clock used all retained proteins, irrespective of tissue specificity. Enrichment ratios and organ assignments for all aptamers are provided in Supplementary Table 6.

### Training organ-specific proteomic clocks

We trained aging clocks for 11 organs (same set as in Oh et al.^4^) on 2,356 samples (1,128 baseline, 1,228 follow-up) using the scikit-learn Python package^43^. For each organ, the preprocessed intensities of organ-enriched aptamers were used to predict chronological age. We employed a nested 5-fold cross-validation scheme to generate unbiased out-of-sample predictions. To prevent data leakage, samples from the same participant were always assigned to the same fold. Each clock consisted of an ensemble of 500 LASSO regression models: for each of the 5 outer training sets 500 bootstrap samples were drawn and a LASSO model was fitted to each. The penalty parameter was tuned within each bootstrap using an inner 5-fold cross-validation to minimize the mean absolute error (MAE). Robust final predictions were obtained by averaging the age predictions across the 500 bootstrap models. Sex-specific clocks were trained using the same pipeline on male and female subsets. Compared with the modeling strategy of Oh et al., our training methodology differed in two minor aspects. First, we did not include sex as a covariate, as sex alone, without protein–sex interaction terms, is not informative for predicting chronological age. Second, we selected the conventional regularization parameter that minimized cross-validated mean absolute error, rather than a more strongly regularized solution that retained 95% of peak cross-validated performance.

### Calculating organ age gaps

Organ age gaps were calculated as the residuals from a LOESS regression of predicted age on chronological age (R function loess, span = 1), fitted separately for each organ, cross-validation fold, and visit. This removed model- and visit-specific systematic differences. Finally, age gaps were standardized within each organ and visit to make them comparable.

### Cross-organ structural equation modelling

To examine whether baseline aging in one organ predicted subsequent aging in others, we used a temporally constrained causal discovery framework that leverages the longitudinal structure of the data. We adapted the structural equation modelling strategy of Tian et al.^19^. Standardized organ age gaps were first sex-adjusted to prevent learning sex-driven dependencies. Graph structure was inferred using the Fast Greedy Equivalence Search (FGES)^44^ implementation in pytetrad (Tetrad v6.8.1: https://github.com/cmu-phil/tetrad), which optimizes the SEM-BIC score (default penalty discount 0.5). Edges were constrained to run only from baseline to follow-up age gaps and only between different organs. The algorithm starts from an empty graph, iteratively adds edges to improve the score, and then performs a backward phase that removes edges until the score can no longer be improved. This procedure was repeated on 500 bootstrap resamples and the consensus network was constructed by retaining edges selected in at least 50 percent of bootstraps. For each retained edge, we estimated its effect size using ordinary least squares regression with the follow-up node as outcome and its baseline parent nodes as predictors. Coefficients were used as edge weights, and p values were FDR-adjusted using the Benjamini–Hochberg procedure with a 5 percent significance threshold. Inclusion in the final network therefore required both reproducibility in the causal search and statistical support in regression.

### Statistical analysis of phenotype associations

Cross-sectional associations between organ age gaps and clinical phenotypes were evaluated using linear mixed-effects models (lmerTest R package^38^). For each organ–phenotype pair, we fitted:

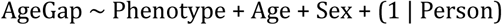

where Person is a random intercept for each participant to account for repeated measurements. Longitudinal associations were evaluated using linear regression models testing the effect of the approximately 10-year change in each phenotype. For each organ–phenotype pair, we fitted:

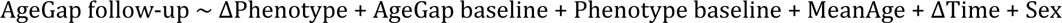

where MeanAge is the participant’s average age across both visits and ΔTime is the time elapsed between baseline and follow-up. In all models, organ age gaps and phenotype variables (baseline, follow-up, and change) were standardized prior to analysis. Missing data for a given phenotype or its longitudinal change led to exclusion from that specific model only, see Extended Table 1 for the distribution and number of missing values per phenotype. P values were adjusted for multiple testing across all organ–phenotype associations using the Benjamini–Hochberg FDR with a 5% significance cut-off.

## Author contributions statement

R.E.N. and T.D.M. conceptualized the study. R.E.N. performed the data exploration, preprocessing, and formal analysis, and wrote the original manuscript, with assistance from M.V.D. and L.C. in data exploration and preprocessing. T.D.M. supervised the project. E.R.R., M.L.D.B and P.S. were responsible for the design and sampling of the Asklepios cohort. J.A.C. oversaw the proteomic data acquisition. All authors contributed to the revision of the manuscript and approved the final version.

## Acknowledgements

This work was supported by the Research Foundation–Flanders (FWO, PhD Fellowship fundamental research grant 11A4425N to R.E.N.) and by FWO research grants G042703 and G083810N for the Asklepios Study. Proteomic measurements were funded by an Investigator-Initiated grant (J.A.C) to the University of Pennsylvania from Bristol Myers Squibb. The funders had no role in the study design; in the collection, analysis, or interpretation of data; in the writing of the manuscript; or in the decision to publish the results.

## Data availability

The individual participant-level data supporting the findings of this study are available for collaborative research upon reasonable request and the execution of appropriate data-sharing agreements; please contact E.R.R. All individual (general and sex-stratified) protein-level regression statistics and aging clock coefficients are provided in the Supplementary Tables.

## Extended Data

**Extended Data Fig. 1:**
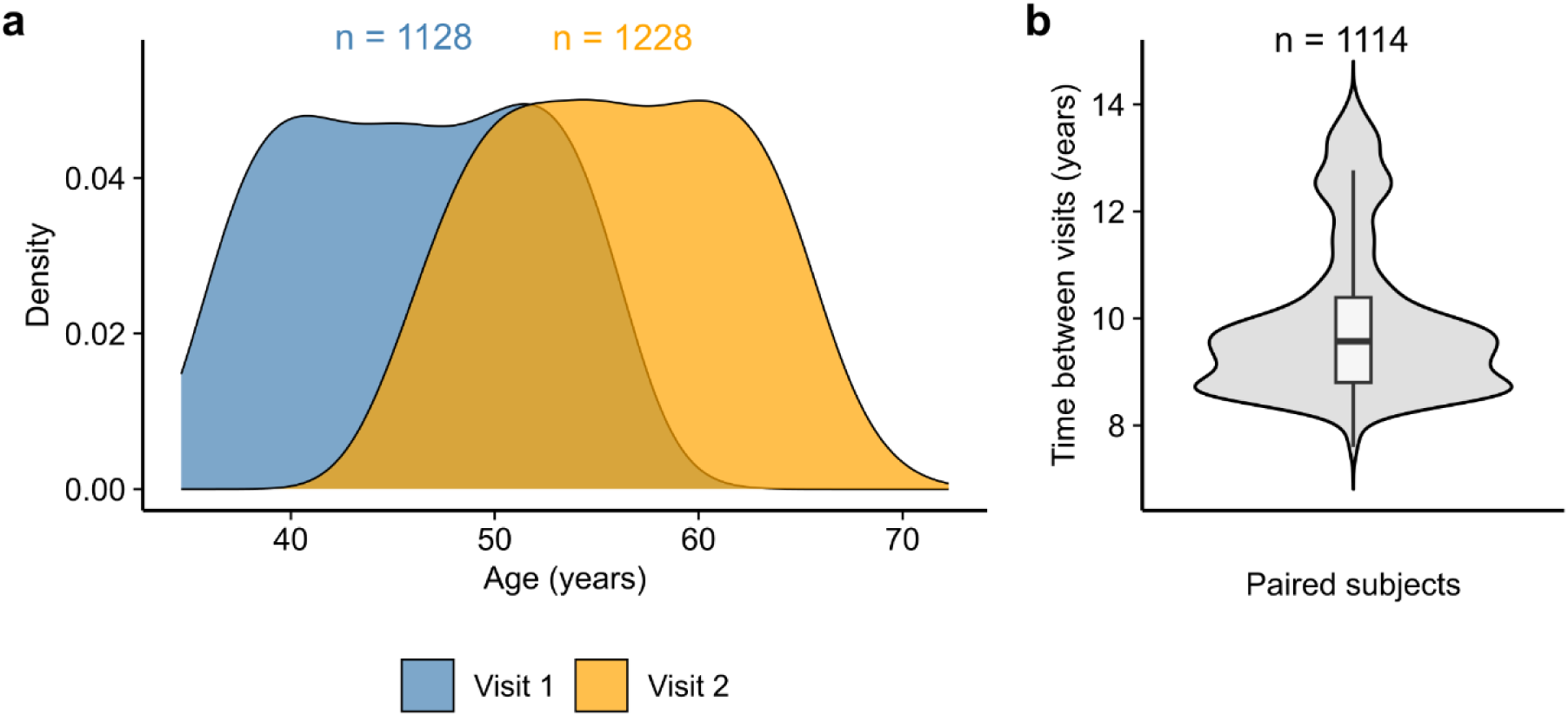
Age distribution and time between visits in the Asklepios cohort. **a**, Age distribution at baseline and follow-up in Asklepios participants with successful proteomic measurements (n = 1,128 at Visit 1; n = 1,228 at Visit 2, out of 1,250 total participants). **b**, Time elapsed between visits for participants with proteomic data at both visits (n = 1,114).

**Extended Data Fig. 2:**
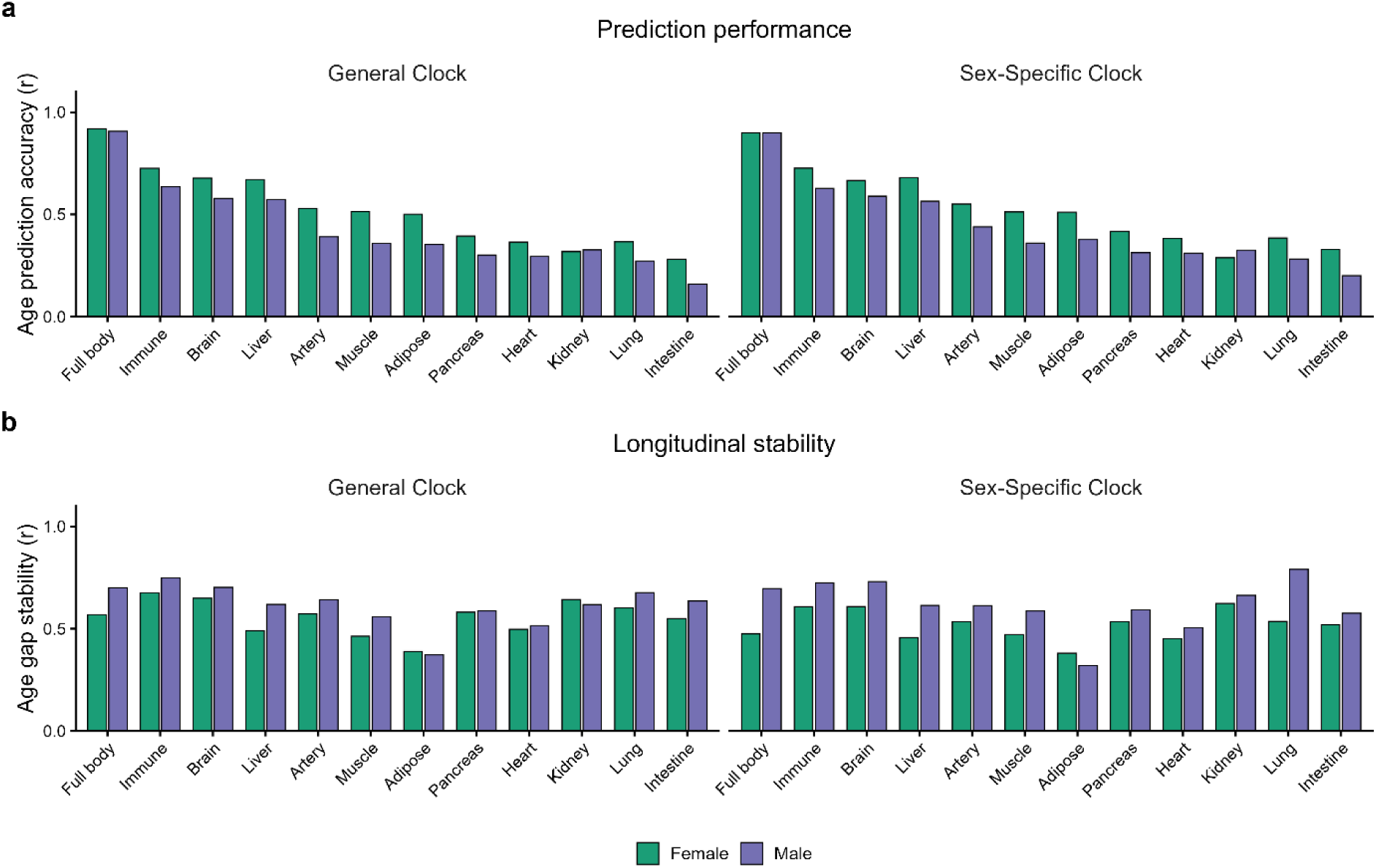
Organ aging clocks achieve consistently higher accuracy in females regardless of training strategy. **a**, Comparison of out-of-fold age-prediction performance (Pearson’s r) between general and sex-specific organ clocks, stratified by sex. **b**, Longitudinal stability of general and sex-specific clocks, calculated as the correlation of z-scored age gaps between baseline and follow-up for each organ, stratified by sex.

**Extended Data Fig. 3:**
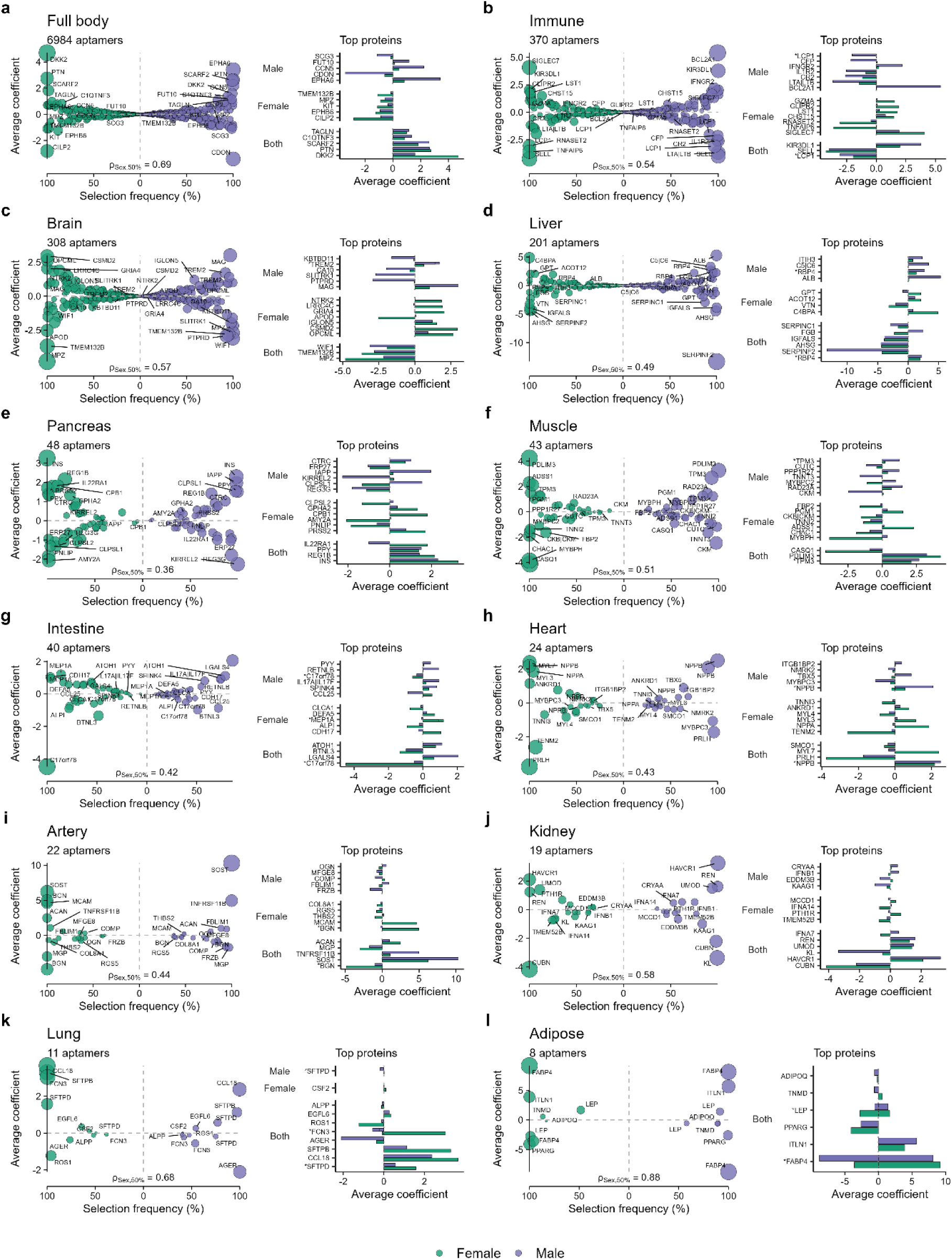
Composition of the sex-specific organ clocks. **a-l**, (left) Average coefficient size and selection frequency of all aptamers across all 500 bootstrapped LASSO models (with 5 folds per iteration), separately per sex. The Spearman correlation (ρSex,50%) was calculated using all aptamers selected in ≥50% of bootstraps in either model. (right) Average coefficient size for the top 10 aptamers in each sex model with corresponding gene symbols. An asterisk (*) denotes gene symbols corresponding to multiple aptamers within the top-10 lists.

**Extended Data Fig. 4:**
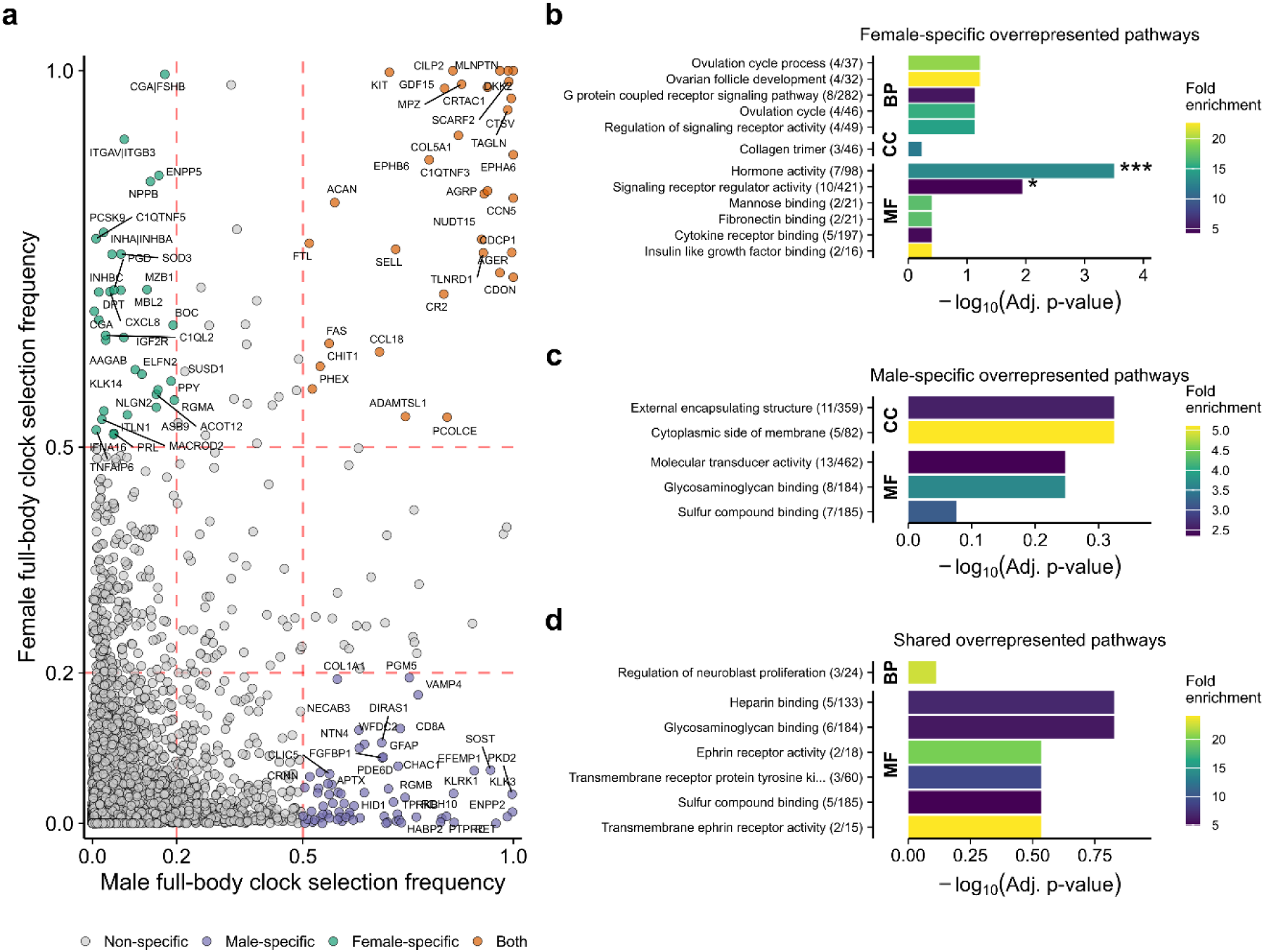
Female-specific full body clock is enriched for hormones. **a**, Selection frequencies of aptamers in the female (y-axis) and male (x-axis) full-body clocks. Selection frequency for each aptamer is defined as the proportion of the bootstrapped LASSO models in which it was selected. Dashed red lines indicate the high (≥0.50) and low (≤0.20) frequency thresholds used to define sex-specific and sex-shared highly selected aptamer lists. These aptamers were converted to unique gene sets via their Entrez gene symbols, resulting in 32 female-specific genes, 69 male-specific genes, and 32 genes frequently selected in both models. **b-d**, Over-Representation Analysis (ORA) of pathways for the gene lists defined in (a). The analysis was performed using the fora function from the fgsea R package^45^, considering pathways with a minimum of 15 and a maximum of 500 genes. Gene sets for Gene Ontology (GO) Biological Process (BP), Cellular Component (CC), Molecular Function (MF), and KEGG pathways were obtained from the Molecular Signatures Database (MSigDB, Homo sapiens) using the msigdbr R package^46^. The background consisted of all genes corresponding to an aptamer that was selected in at least one bootstrapped LASSO (n = 5,785 genes). The plots show up to the top five most significantly enriched pathways per database with an FDR < 1. FDR correction applied separately for each pathway group and gene list. Significance is denoted by asterisks (*Adj. p < 0.05, **Adj. p < 0.01, ***Adj. p < 0.001). No KEGG pathways were significantly enriched in any of the analyses.

**Extended Data Fig. 5:**
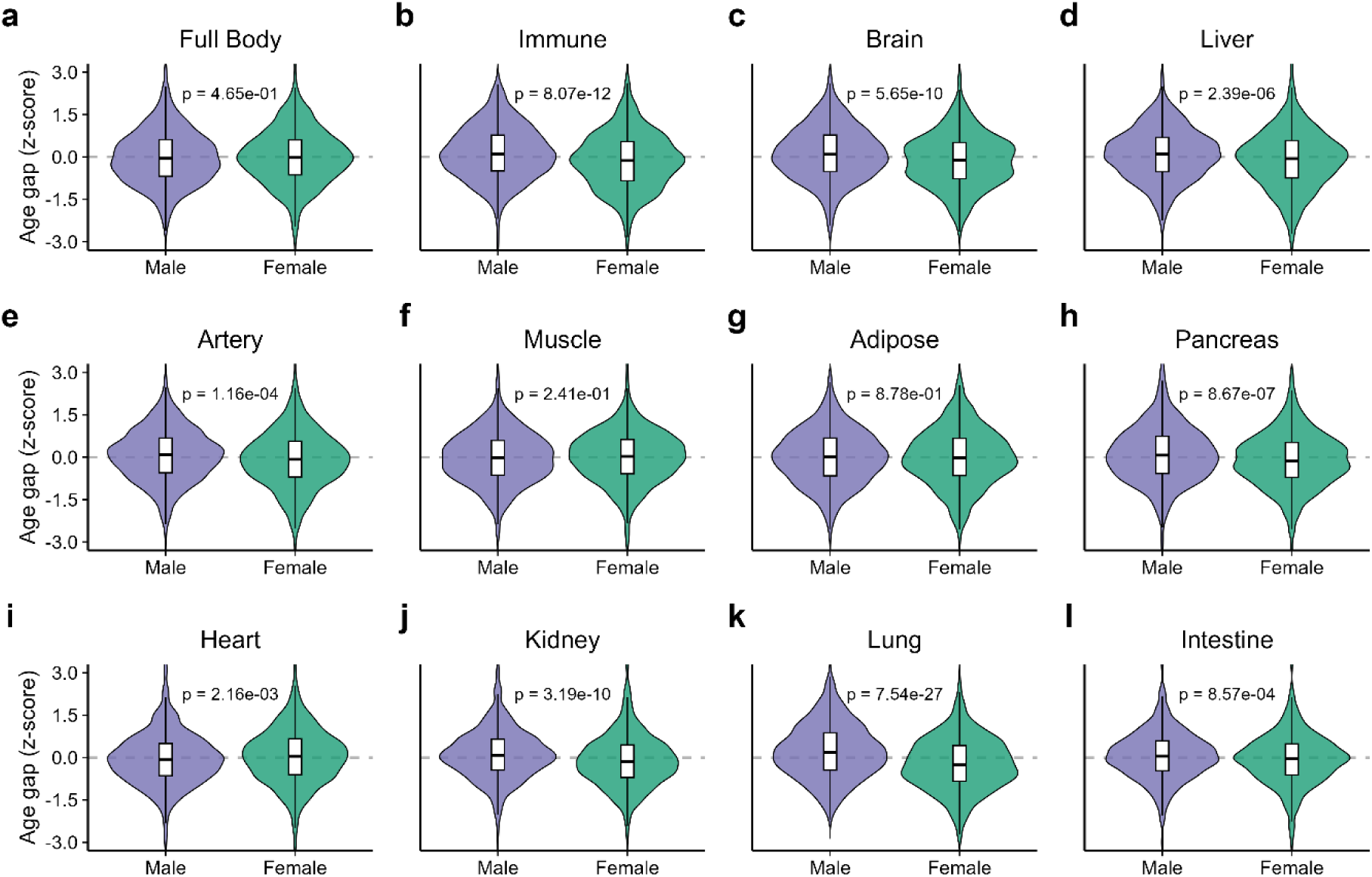
Male organs show accelerated aging for most organs in the general clocks. **a-l**, Comparison of standardized age gaps from the general sex-agnostic aging clocks between males and females for each of the 11 organs (b-l) and for the full body (a). P-values are obtained from a two-sided Wilcoxon rank-sum test. Organs are ordered according to the overall age prediction performance.

**Extended Data Fig. 6:**
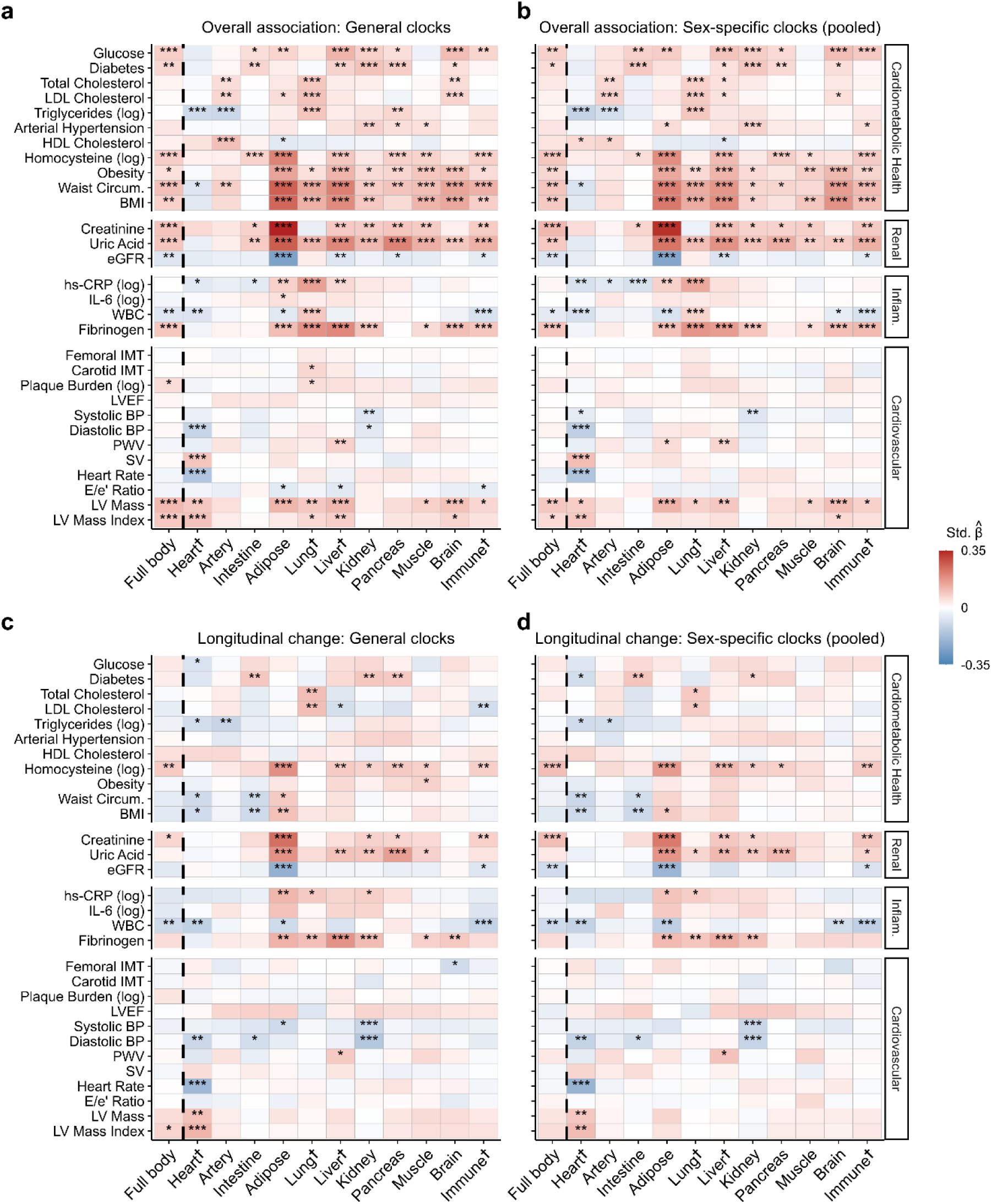
Phenotype associations from sex-specific clocks largely align with results from general clock. **a–b**, Cross-sectional associations between organ age gaps and cardiometabolic, renal, inflammatory, and cardiovascular markers across all samples (n = 2,356; 1,128 baseline and 1,228 follow-up). Panel a uses age gaps from the general organ clocks; panel b uses age gaps from sex-specific clocks (pooled across sexes). Linear mixed-effects models regressed z-scored organ age gaps on z-scored clinical markers, adjusted for age and sex with a random intercept for participant. **c–d**, Longitudinal associations between (approximately) 10-year changes in organ age gaps and changes in the same markers (n = 1,114 individuals with proteomic data at both visits). Panel c uses general clocks; panel d uses sex-specific clocks (pooled). Linear models predicted follow-up z-scored organ age gaps from z-scored biomarker changes, adjusted for z-scored baseline age gap, z-scored baseline biomarker level, mean age, age change, and sex. P values were FDR-adjusted within each analysis (*Adj. p < 0.05, **Adj. p < 0.01, ***Adj. p < 0.001). **†**Organs with clocks that have strong independent validation for uniquely predicting diseases specific to their respective organ^10^. Abbreviations: BMI, Body Mass Index; BP, Blood Pressure; eGFR, estimated Glomerular Filtration Rate; hs-CRP, high-sensitivity C-reactive protein; IL-6, Interleukin-6; IMT, Intima-Media Thickness; LV, Left Ventricular; LVEF, Left Ventricular Ejection Fraction; NSAID, Non-Steroidal Anti-Inflammatory Drug; PWV, Pulse Wave Velocity; RAS, Renin–Angiotensin System; SV, Stroke Volume; WBC, White Blood Cell count.

**Extended Data Fig. 7.**
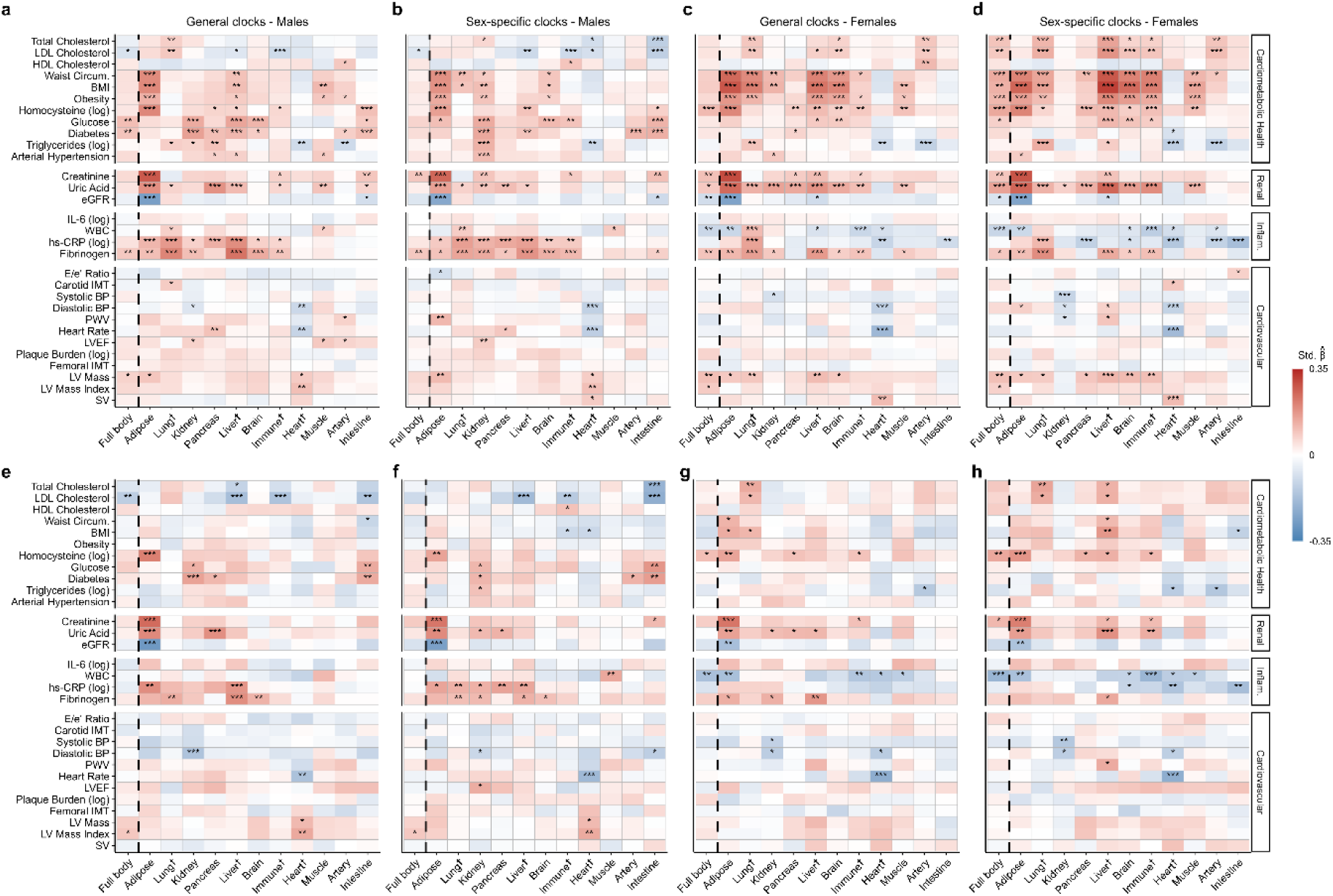
Sex-stratified associations between organ age gaps and clinical phenotypes using general versus sex-specific clocks. **a–d**, Cross-sectional associations between organ age gaps and cardiometabolic, renal, inflammatory, and cardiovascular markers in males (a–b; n = 1,156 samples, 555 baseline and 601 follow-up) and females (c–d; n = 1,200 samples, 573 baseline and 627 follow-up). Panels a and c use age gaps from the general organ clocks; panels b and d use age gaps from the sex-specific clocks. Linear mixed-effects models regressed z-scored organ age gaps on sex-stratified z-scored clinical markers, adjusted for age with a random intercept for participant. **e–h**, Longitudinal associations between (approximately) 10-year changes in organ age gaps and corresponding changes in the same markers in males (e–f; n = 547 individuals) and females (g–h; n = 567 individuals) with proteomic data at both visits. Panels e and g use general clocks; panels f and h use sex-specific clocks. Linear models predicted follow-up z-scored organ age gaps from sex-stratified z-scored biomarker changes, adjusted for z-scored baseline age gap, sex-stratified z-scored baseline biomarker level, mean age, and age change. P values were FDR-adjusted separately for each of the eight analyses (*Adj. p < 0.05, **Adj. p < 0.01, ***Adj. p < 0.001). **†**Organs with clocks that have strong independent validation for uniquely predicting diseases specific to their respective organ^10^. Abbreviations: BMI, Body Mass Index; BP, Blood Pressure; eGFR, estimated Glomerular Filtration Rate; hs-CRP, high-sensitivity C-reactive protein; IL-6, Interleukin-6; IMT, Intima-Media Thickness; LV, Left Ventricular; LVEF, Left Ventricular Ejection Fraction; NSAID, Non-Steroidal Anti-Inflammatory Drug; PWV, Pulse Wave Velocity; RAS, Renin–Angiotensin System; SV, Stroke Volume; WBC, White Blood Cell count.

**Extended Data Fig. 8:**
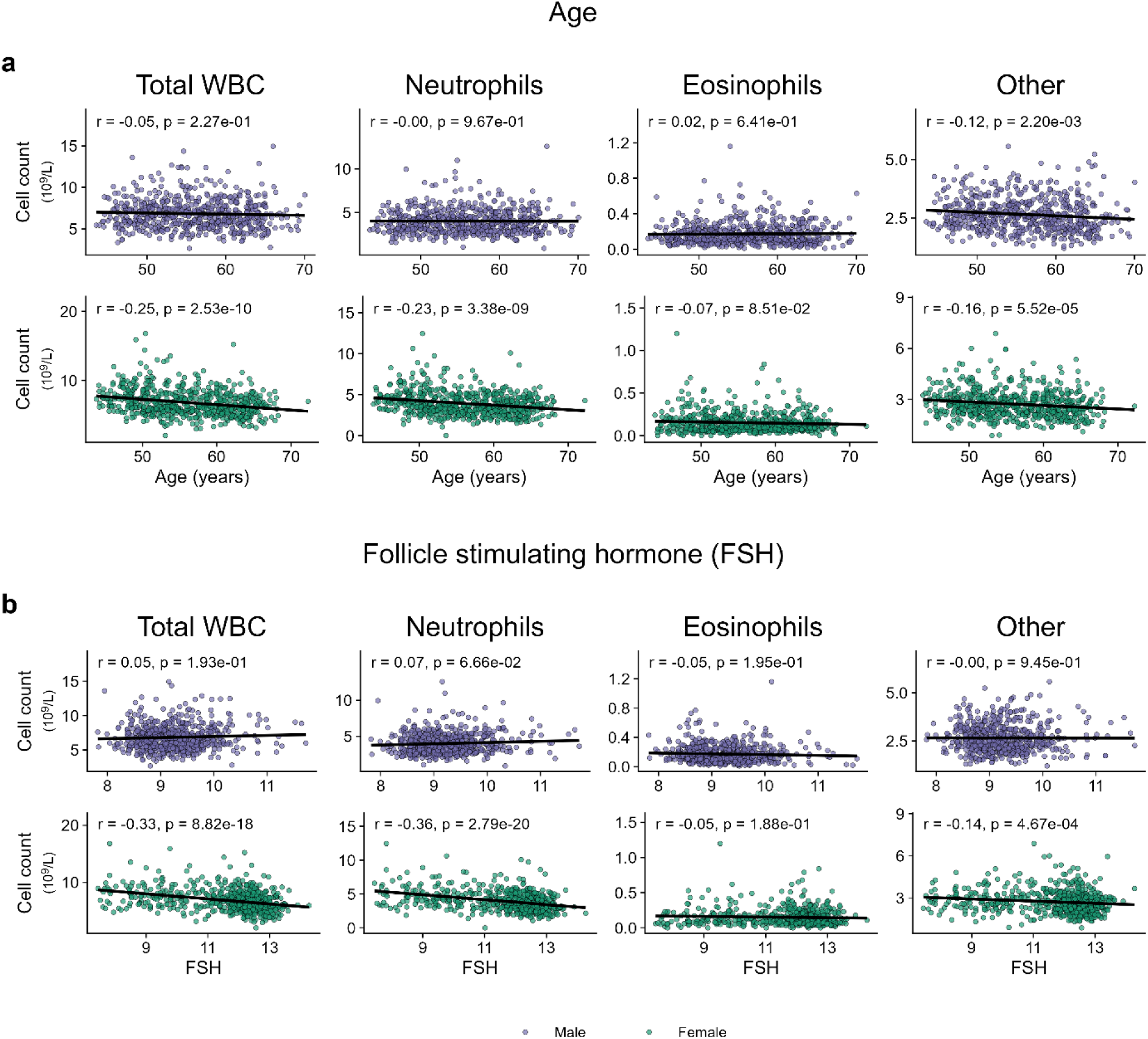
Sex-specific relationships of white blood cell subtypes with age and follicle-stimulating hormone. **a,** Correlations between white blood cell (WBC) counts and age. **b,** Correlations between white blood cell (WBC) counts and follicle stimulating hormone (FSH) concentrations. Pearson’s r and unadjusted p-values are shown. The analysis used data from the follow-up visit for participants with proteomic clock predictions (n = 1,221, 3 NAs).

**Extended Data Table 1:**
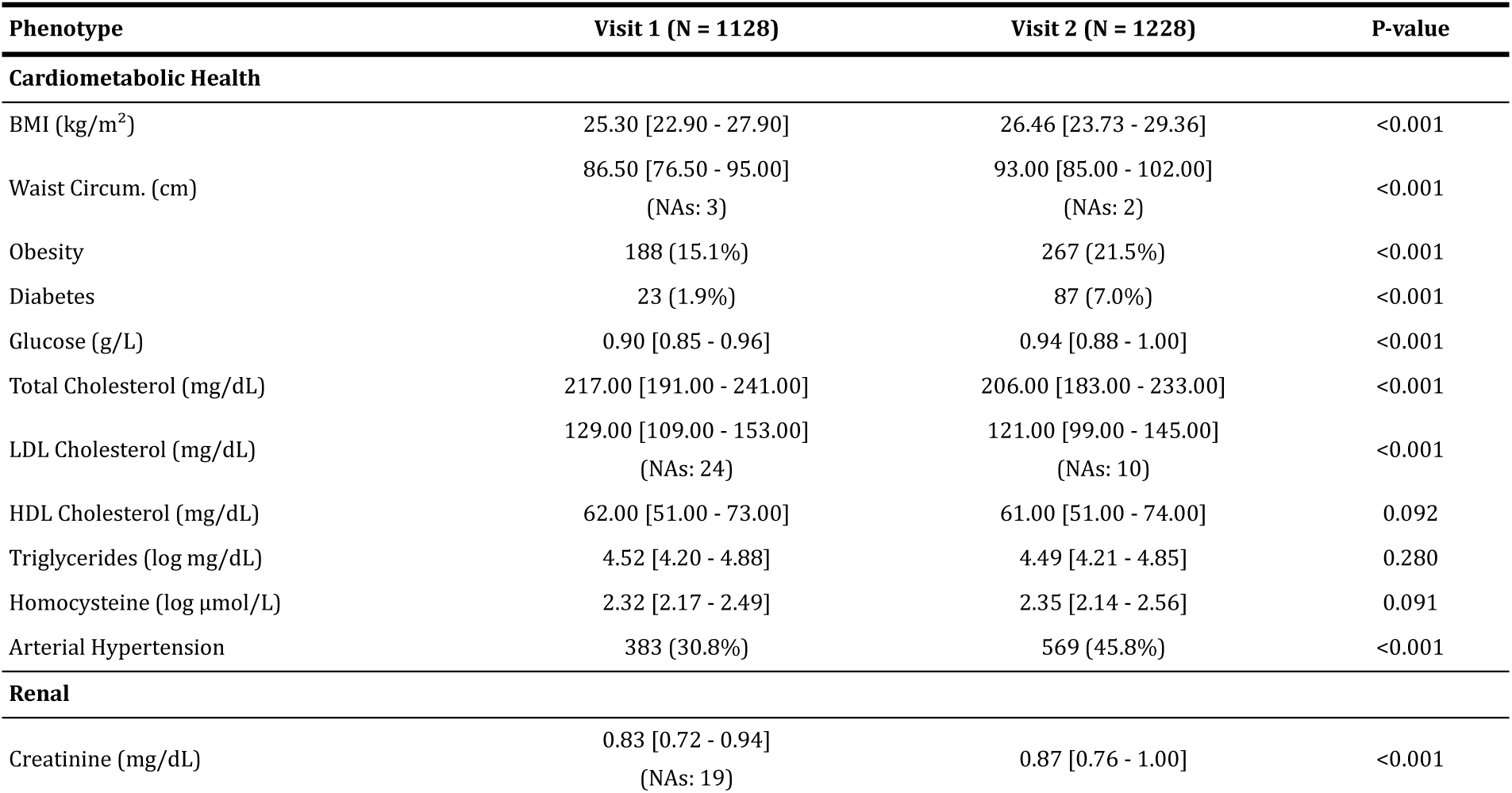
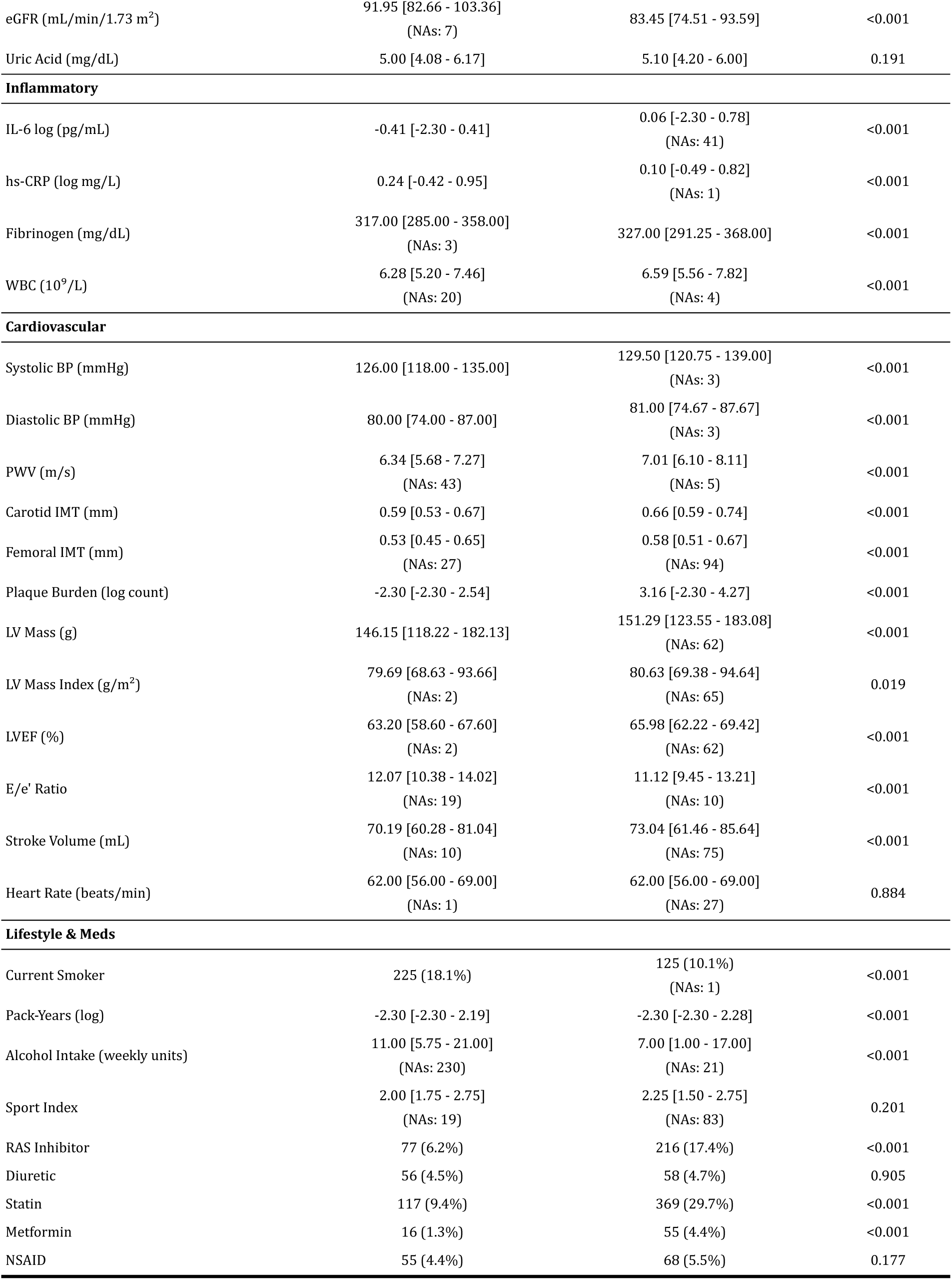
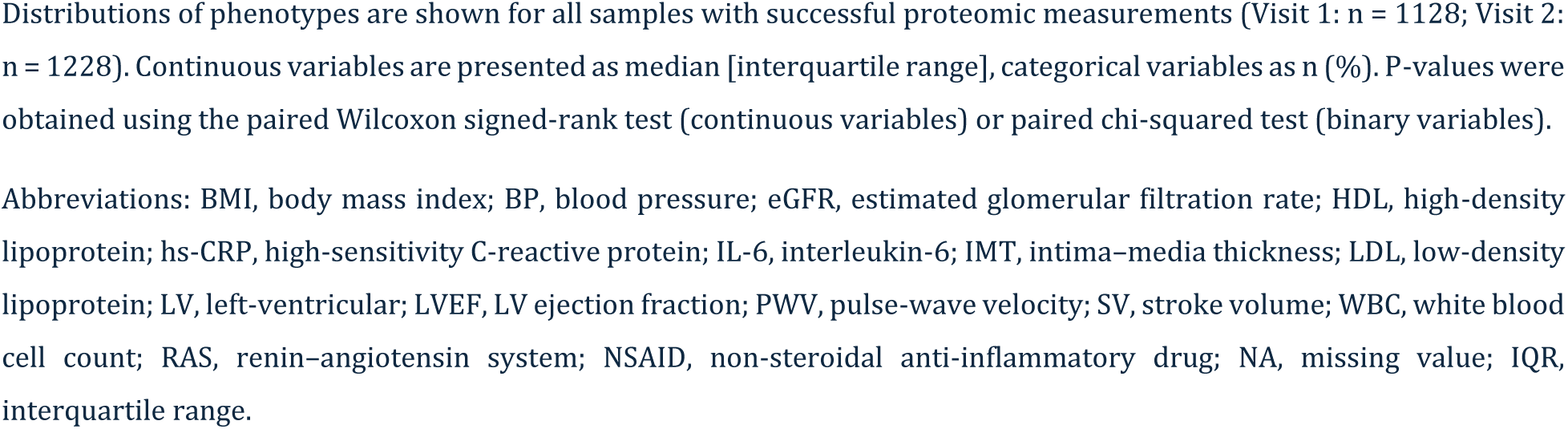
Distributions of clinical phenotypes at baseline and follow-up in the Asklepios cohort.

| Phenotype | Visit 1 (N = 1128) | Visit 2 (N = 1228) | P-value |
| --- | --- | --- | --- |
| **Cardiometabolic Health** |  |  |  |
| BMI (kg/m<sup>2</sup>) | 25.30 [22.90 - 27.90] | 26.46 [23.73 - 29.36] | <0.001 |
| Waist Circum. (cm) | 86.50 [76.50 - 95.00]<br>(NAs: 3) | 93.00 [85.00 - 102.00]<br>(NAs: 2) | <0.001 |
| Obesity | 188 (15.1%) | 267 (21.5%) | <0.001 |
| Diabetes | 23 (1.9%) | 87 (7.0%) | <0.001 |
| Glucose (g/L) | 0.90 [0.85 - 0.96] | 0.94 [0.88 - 1.00] | <0.001 |
| Total Cholesterol (mg/dL) | 217.00 [191.00 - 241.00] | 206.00 [183.00 - 233.00] | <0.001 |
| LDL Cholesterol (mg/dL) | 129.00 [109.00 - 153.00]<br>(NAs: 24) | 121.00 [99.00 - 145.00]<br>(NAs: 10) | <0.001 |
| HDL Cholesterol (mg/dL) | 62.00 [51.00 - 73.00] | 61.00 [51.00 - 74.00] | 0.092 |
| Triglycerides (log mg/dL) | 4.52 [4.20 - 4.88] | 4.49 [4.21 - 4.85] | 0.280 |
| Homocysteine (log μmol/L) | 2.32 [2.17 - 2.49] | 2.35 [2.14 - 2.56] | 0.091 |
| Arterial Hypertension | 383 (30.8%) | 569 (45.8%) | <0.001 |
| **Renal** |  |  |  |
| Creatinine (mg/dL) | 0.83 [0.72 - 0.94]<br>(NAs: 19) | 0.87 [0.76 - 1.00] | <0.001 |
| eGFR (mL/min/1.73 m<sup>2</sup>) | 91.95 [82.66 - 103.36]<br>(NAs: 7) | 83.45 [74.51 - 93.59] | <0.001 |
| Uric Acid (mg/dL) | 5.00 [4.08 - 6.17] | 5.10 [4.20 - 6.00] | 0.191 |
| **Inflammatory** |  |  |  |
| IL-6 log (pg/mL) | -0.41 [-2.30 - 0.41] | 0.06 [-2.30 - 0.78]<br>(NAs: 41) | <0.001 |
| hs-CRP (log mg/L) | 0.24 [-0.42 - 0.95] | 0.10 [-0.49 - 0.82]<br>(NAs: 1) | <0.001 |
| Fibrinogen (mg/dL) | 317.00 [285.00 - 358.00]<br>(NAs: 3) | 327.00 [291.25 - 368.00] | <0.001 |
| WBC (10<sup>9</sup>/L) | 6.28 [5.20 - 7.46]<br>(NAs: 20) | 6.59 [5.56 - 7.82]<br>(NAs: 4) | <0.001 |
| **Cardiovascular** |  |  |  |
| Systolic BP (mmHg) | 126.00 [118.00 - 135.00] | 129.50 [120.75 - 139.00]<br>(NAs: 3) | <0.001 |
| Diastolic BP (mmHg) | 80.00 [74.00 - 87.00] | 81.00 [74.67 - 87.67]<br>(NAs: 3) | <0.001 |
| PWV (m/s) | 6.34 [5.68 - 7.27]<br>(NAs: 43) | 7.01 [6.10 - 8.11]<br>(NAs: 5) | <0.001 |
| Carotid IMT (mm) | 0.59 [0.53 - 0.67] | 0.66 [0.59 - 0.74] | <0.001 |
| Femoral IMT (mm) | 0.53 [0.45 - 0.65]<br>(NAs: 27) | 0.58 [0.51 - 0.67]<br>(NAs: 94) | <0.001 |
| Plaque Burden (log count) | -2.30 [-2.30 - 2.54] | 3.16 [-2.30 - 4.27] | <0.001 |
| LV Mass (g) | 146.15 [118.22 - 182.13] | 151.29 [123.55 - 183.08]<br>(NAs: 62) | <0.001 |
| LV Mass Index (g/m<sup>2</sup>) | 79.69 [68.63 - 93.66]<br>(NAs: 2) | 80.63 [69.38 - 94.64]<br>(NAs: 65) | 0.019 |
| LVEF (%) | 63.20 [58.60 - 67.60]<br>(NAs: 2) | 65.98 [62.22 - 69.42]<br>(NAs: 62) | <0.001 |
| E/e' Ratio | 12.07 [10.38 - 14.02]<br>(NAs: 19) | 11.12 [9.45 - 13.21]<br>(NAs: 10) | <0.001 |
| Stroke Volume (mL) | 70.19 [60.28 - 81.04]<br>(NAs: 10) | 73.04 [61.46 - 85.64]<br>(NAs: 75) | <0.001 |
| Heart Rate (beats/min) | 62.00 [56.00 - 69.00]<br>(NAs: 1) | 62.00 [56.00 - 69.00]<br>(NAs: 27) | 0.884 |
| **Lifestyle & Meds** |  |  |  |
| Current Smoker | 225 (18.1%) | 125 (10.1%)<br>(NAs: 1) | <0.001 |
| Pack-Years (log) | -2.30 [-2.30 - 2.19] | -2.30 [-2.30 - 2.28] | <0.001 |
| Alcohol Intake (weekly units) | 11.00 [5.75 - 21.00]<br>(NAs: 230) | 7.00 [1.00 - 17.00]<br>(NAs: 21) | <0.001 |
| Sport Index | 2.00 [1.75 - 2.75]<br>(NAs: 19) | 2.25 [1.50 - 2.75]<br>(NAs: 83) | 0.201 |
| RAS Inhibitor | 77 (6.2%) | 216 (17.4%) | <0.001 |
| Diuretic | 56 (4.5%) | 58 (4.7%) | 0.905 |
| Statin | 117 (9.4%) | 369 (29.7%) | <0.001 |
| Metformin | 16 (1.3%) | 55 (4.4%) | <0.001 |
| NSAID | 55 (4.4%) | 68 (5.5%) | 0.177 |

Distributions of phenotypes are shown for all samples with successful proteomic measurements (Visit 1: n = 1128; Visit 2: n = 1228). Continuous variables are presented as median [interquartile range], categorical variables as n (%). P-values were obtained using the paired Wilcoxon signed-rank test (continuous variables) or paired chi-squared test (binary variables).
Abbreviations: BMI, body mass index; BP, blood pressure; eGFR, estimated glomerular filtration rate; HDL, high-density lipoprotein; hs-CRP, high-sensitivity C-reactive protein; IL-6, interleukin-6; IMT, intima-media thickness; LDL, low-density lipoprotein; LV, left-ventricular; LVEF, LV ejection fraction; PWV, pulse-wave velocity; SV, stroke volume; WBC, white blood cell count; RAS, renin-angiotensin system; NSAID, non-steroidal anti-inflammatory drug; NA, missing value; IQR, interquartile range.

## References

1. Lo pez-Otín, C., Blasco, M. A., Partridge, L., Serrano, M. & Kroemer, G. The Hallmarks of Aging. Cell 153, 1194–1217 (2013).

2. Liu, Z. et al. A new aging measure captures morbidity and mortality risk across diverse subpopulations from NHANES IV: A cohort study. PLOS Med. 15, e1002718 (2018).

3. Schaum, N. et al. Aging hallmarks exhibit organ-specific temporal signatures. Nature 583, 596–602 (2020).

4. Oh, H. S.-H. et al. Organ aging signatures in the plasma proteome track health and disease. Nature 624, 164–172 (2023).

5. Tanaka, T. et al. Plasma proteomic signature of age in healthy humans. Aging Cell 17, e12799 (2018).

6. Hannum, G. et al. Genome-wide methylation profiles reveal quantitative views of human aging rates. Mol. Cell 49, 359–367 (2013).

7. Peters, M. J. et al. The transcriptional landscape of age in human peripheral blood. Nat. Commun. 6, 8570 (2015).

8. Galkin, F. et al. Human Gut Microbiome Aging Clock Based on Taxonomic Profiling and Deep Learning. iScience 23, 101199 (2020).

9. Goeminne, L. J. E. et al. Plasma protein-based organ-specific aging and mortality models unveil diseases as accelerated aging of organismal systems. Cell Metab. 37, 205–222.e6 (2025).

10. Kivima ki, M. et al. Proteomic organ-specific ageing signatures and 20-year risk of age-related diseases: the Whitehall II observational cohort study. Lancet Digit. Health 7, e195–e204 (2025).

11. Tang, J. et al. Longitudinal serum proteome mapping reveals biomarkers for healthy ageing and related cardiometabolic diseases. Nat. Metab. 7, 166–181 (2025).

12. Lehallier, B. et al. Undulating changes in human plasma proteome profiles across the lifespan. Nat. Med. 25, 1843–1850 (2019).

13. Marijic, J. et al. Decreased expression of voltage- and Ca(2+)-activated K(+) channels in coronary smooth muscle during aging. Circ. Res. 88, 210–216 (2001).

14. de Lores Arnaiz, G. R. & Ordieres, M. G. L. Brain Na(+), K(+)-ATPase Activity In Aging and Disease. Int. J. Biomed. Sci. IJBS 10, 85–102 (2014).

15. Ostan, R. et al. Gender, aging and longevity in humans: an update of an intriguing/neglected scenario paving the way to a gender-specific medicine. Clin. Sci. 130, 1711–1725 (2016).

16. Sathyan, S. et al. Plasma proteomic profile of age, health span, and all-cause mortality in older adults. Aging Cell 19, e13250 (2020).

17. Liu, F. et al. Mid-life plasma proteins associated with late-life prefrailty and frailty: a proteomic analysis. GeroScience 46, 5247–5265 (2024).

18. The GTEx Consortium. The GTEx Consortium atlas of genetic regulatory effects across human tissues. Science 369, 1318–1330 (2020).

19. Tian, Y. E. et al. Heterogeneous aging across multiple organ systems and prediction of chronic disease and mortality. Nat. Med. 29, 1221–1231 (2023).

20. Sengene s, C., Berlan, M., De Glisezinski, I., Lafontan, M. & Galitzky, J. Natriuretic peptides: a new lipolytic pathway in human adipocytes. FASEB J. Off. Publ. Fed. Am. Soc. Exp. Biol. 14, 1345–1351 (2000).

21. Bordicchia, M. et al. Cardiac natriuretic peptides act via p38 MAPK to induce the brown fat thermogenic program in mouse and human adipocytes. J. Clin. Invest. 122, 1022–1036 (2012).

22. Austad, S. N. & Fischer, K. E. Sex Differences in Lifespan. Cell Metab. 23, 1022–1033 (2016).

23. Chen, Y. et al. Difference in Leukocyte Composition between Women before and after Menopausal Age, and Distinct Sexual Dimorphism. PloS One 11, e0162953 (2016).

24. Wang, Y. et al. Organ-specific proteomic aging clocks predict disease and longevity across diverse populations. Nat. Aging 1–19 (2025) doi:10.1038/s43587-025-01016-8.

25. Bomback, A. S. & Toto, R. Dual Blockade of the Renin–Angiotensin–Aldosterone System: Beyond the ACE Inhibitor and Angiotensin-II Receptor Blocker Combination. Am. J. Hypertens. 22, 1032–1040 (2009).

26. Bohn, B. et al. A Proteomic Approach for Investigating the Pleiotropic Effects of Statins in the Atherosclerosis Risk in Communities (ARIC) Study. J. Proteomics 272, 104788 (2023).

27. Franceschi, C., Garagnani, P., Parini, P., Giuliani, C. & Santoro, A. Inflammaging: a new immune-metabolic viewpoint for age-related diseases. Nat. Rev. Endocrinol. 14, 576–590 (2018).

28. Coelho, M., Oliveira, T. & Fernandes, R. Biochemistry of adipose tissue: an endocrine organ. Arch. Med. Sci. AMS 9, 191–200 (2013).

29. Liu, Y. & Li, C. Hormone Therapy and Biological Aging in Postmenopausal Women. JAMA Netw. Open 7, e2430839 (2024).

30. Field, A. E. et al. DNA Methylation Clocks in Aging: Categories, Causes, and Consequences. Mol. Cell 71, 882–895 (2018).

31. Bell, C. G. et al. DNA methylation aging clocks: challenges and recommendations. Genome Biol. 20, 249 (2019).

32. Fitzgerald, K. N. et al. Potential reversal of epigenetic age using a diet and lifestyle intervention: a pilot randomized clinical trial. Aging 13, 9419–9432 (2021).

33. Zhang, Q. et al. Improved precision of epigenetic clock estimates across tissues and its implication for biological ageing. Genome Med. 11, 54 (2019).

34. Veerapaneni, V. V. et al. Circulating Secretoglobin Family 1A Member 1 (SCGB1A1) Levels as a Marker of Biomass Smoke Induced Chronic Obstructive Pulmonary Disease. Toxics 9, 208 (2021).

35. Rietzschel, E.-R. et al. Rationale, design, methods and baseline characteristics of the Asklepios Study. Eur. J. Cardiovasc. Prev. Rehabil. Off. J. Eur. Soc. Cardiol. Work. Groups Epidemiol. Prev. Card. Rehabil. Exerc. Physiol. 14, 179–191 (2007).

36. Dib, M.-J. et al. Proteome-Wide Genetic Investigation of Large Artery Stiffness. JACC Basic Transl. Sci. 9, 1178–1191 (2024).

37. Curran, P. J. & Bauer, D. J. The Disaggregation of Within-Person and Between-Person Effects in Longitudinal Models of Change. Annu. Rev. Psychol. 62, 583–619 (2011).

38. Kuznetsova, A., Brockhoff, P. B. & Christensen, R. H. B. lmerTest Package: Tests in Linear Mixed Effects Models. J. Stat. Softw. 82, 1–26 (2017).

39. Korotkevich, G., Sukhov, V. & Sergushichev, A. Fast gene set enrichment analysis. 060012 Preprint at 10.1101/060012 (2019).

40. Liberzon, A. et al. The Molecular Signatures Database (MSigDB) hallmark gene set collection. Cell Syst. 1, 417–425 (2015).

41. Robinson, M. D. & Oshlack, A. A scaling normalization method for differential expression analysis of RNA-seq data. Genome Biol. 11, R25 (2010).

42. Candia, J., Daya, G. N., Tanaka, T., Ferrucci, L. & Walker, K. A. Assessment of variability in the plasma 7k SomaScan proteomics assay. Sci. Rep. 12, 17147 (2022).

43. Pedregosa, F. et al. Scikit-learn: Machine Learning in Python. J. Mach. Learn. Res. 12, 2825–2830 (2011).

44. Ramsey, J., Glymour, M., Sanchez-Romero, R. & Glymour, C. A million variables and more: the Fast Greedy Equivalence Search algorithm for learning high-dimensional graphical causal models, with an application to functional magnetic resonance images. Int. J. Data Sci. Anal. 3, 121–129 (2017).

45. fgsea. Bioconductor http://bioconductor.org/packages/fgsea/.

46. msigdb. Bioconductor http://bioconductor.org/packages/msigdb/.

