## Supplementary Methods Note for "Longitudinal dynamics of organ-specific proteomic aging clocks over a decade of midlife"

#### *Normalization of raw SomaScan data*

Raw SomaScan v4.1 7k .adat files from both Asklepios rounds were jointly normalized using the internal-reference pipeline of Candia et al.[1], implemented via the authors' R script (https://osf.io/srgef). Although the proteomic measurements from the two visits were performed about one year apart, they were normalized together because both used the same v4.1 plasma calibrator material and hybridization-control reagents. The pipeline sequentially performs (i) hybridization-control normalization, (ii) median signal normalization on calibrators, (iii) plate-scale normalization, (iv) SOMAmer-specific inter-plate calibration and (v) median-signal normalization across all sample types. Note that this pipeline removes assay-technical plate and batch effects but does not correct technical differences arising before the SomaScan measurements (e.g. storage-time artefacts), which are addressed further below.

#### *Aptamer filtering*

Of the 7,523 aptamers measured by the SomaScan 7K platform, only the 7,289 targeting human proteins were retained. Aptamers for non-human proteins, hybridization controls, and other technical probes were excluded. These 7,289 aptamers quantified 6,402 unique proteins (based on UniProt identifiers). Detailed annotations for each aptamer are provided in Supplementary Table S8. Protein intensities were measured in relative fluorescent units and log2-transformed to make distributions more normal-like.

#### *Sample filtering*

Two samples (one from Round 1 and one from Round 2) with strongly deviating intensity distributions were removed (Supplementary Methods Note Fig. 1a). In addition, 22 Round 1 samples formed a completely separate cluster in a UMAP analysis (Supplementary Methods Note Fig. 1b). This cluster could not be explained by any biological variable, and all affected samples were collected within a short time window, suggesting a local technical batch effect. These samples were therefore excluded from all analyses. Finally, three participants with outlying ages (29, 75 and 75 years at follow-up) were removed (both rounds) to avoid disproportionate leverage on aging model coefficients.

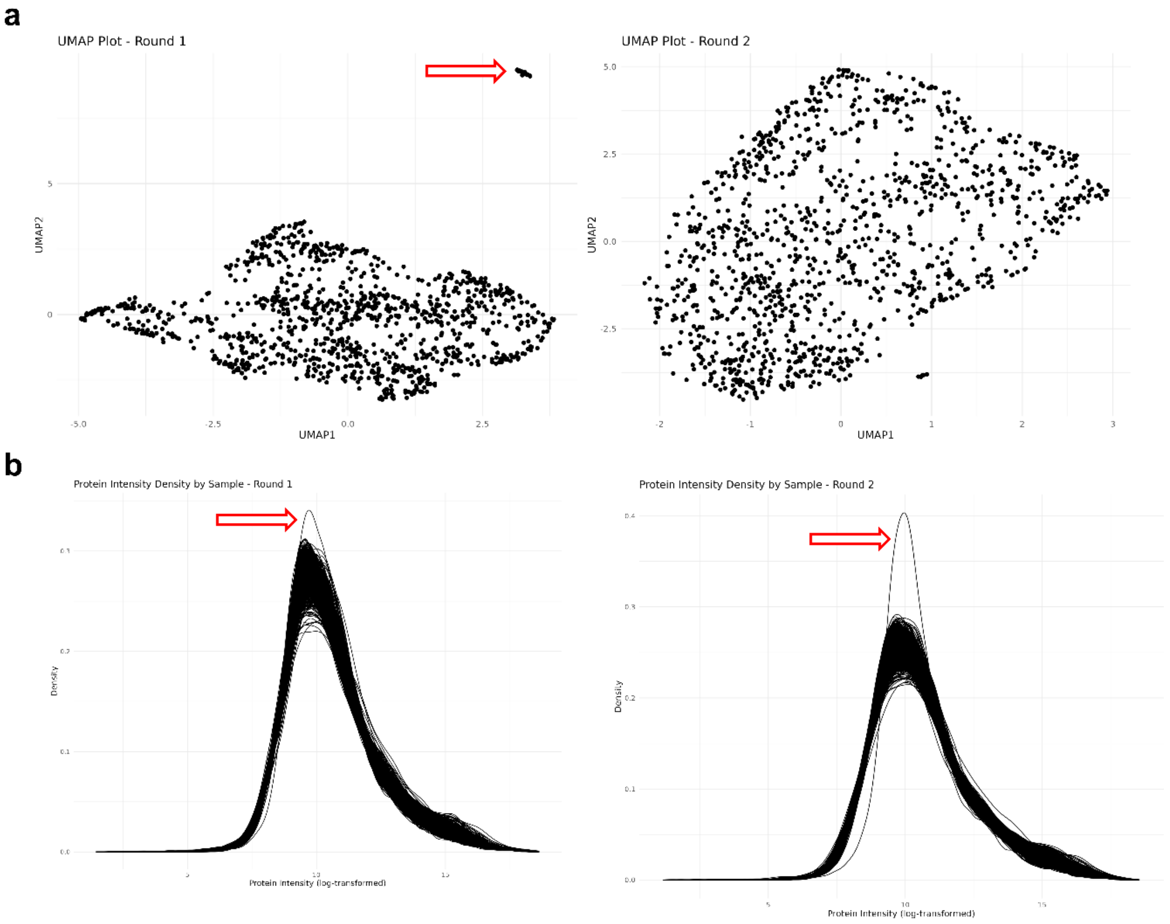


*Supplementary Methods Note Fig. 1: Filtering of outlier samples.*

#### *Addressing remaining technical variability and batch effects*

We identified three sources of technical variation: (1) a general batch effect between baseline and follow-up, likely related to extended storage (~10 years) and non-simultaneous processing (~one year apart), (2) a storage-time drift, and (3) batch effects across plates run on different days.

We first corrected (2) and (3) within each visit (Supplementary Methods Note Fig. 2a–d). Storage-time effects were removed by fitting robust linear models with rlm from the MASS package[2] in R, regressing each aptamer's intensity on the number of days passed since the first sample was collected, separately per plate-run date. Adjusted values were obtained as the model residuals plus the original mean intensity of that aptamer within the same plate-run date. The resulting values were then harmonized using the empirical Bayes ComBat method[3] (implemented in the sva[4] R package), which corrects systematic plate-run differences in mean and variance. Both issues were generally modest and affected only a small minority of aptamers.

Finally, the between-visit batch effect (1) was removed by aligning baseline to follow-up via per-aptamer location and scale adjustments. To avoid eliminating true biological differences arising from the ~10-year age shift, the parameters were estimated in a balanced subset of 561 participants (mean age 51.1 years, 49.7 percent women) matched across visits by sex and one-year age bins (Supplementary Methods Note Fig. 2e–g). These estimated parameters were then applied to the full dataset.

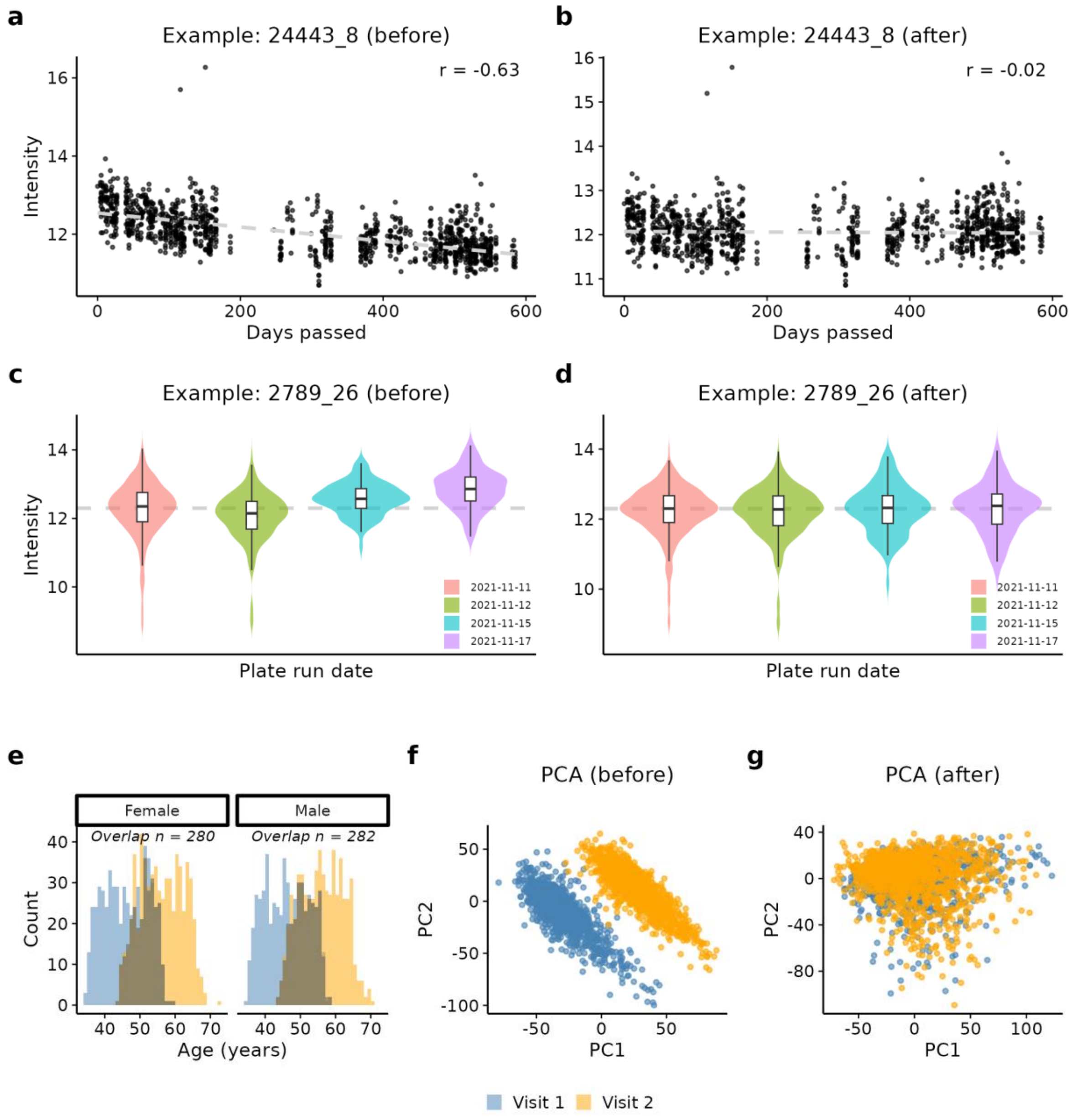


*Supplementary Methods Note Fig. 2: Removing technical artifacts in longitudinal proteomic data*

**a–d**, Illustration of the correction of remaining technical variation using aptamers selected for displaying strong batch effects in the baseline visit. **a–b**, Signal intensity versus days since first sample collection for aptamer 24443_8 before (a) and after (b) robust linear model regression. **c–d**, Intensity distributions across plate-run dates for aptamer 2789_26 before (c) and after (d) ComBat correction harmonizing location and scale. **e**, The dark grey regions show the overlapping reference subset (n = 280 female + 282 male) matched on one-year age bins and sex across visits. In this subset, we estimated the location and scale factors used to correct between-visit batch effect related differences while preserving those driven by the age shift. **f–g**, Principal Component Analysis (PCA) of the full dataset before (f) and after (g) normalization using parameters derived from this reference subset.

### References

1. Candia, J., Daya, G. N., Tanaka, T., Ferrucci, L. & Walker, K. A. Assessment of variability in the plasma 7k SomaScan proteomics assay. Sci. Rep. **12**, 17147 (2022).

2. Venables, W. N. & Ripley, B. D. Modern Applied Statistics with S 4th edn (Springer, 2002).

3. Johnson, W. E., Li, C. & Rabinovic, A. Adjusting batch effects in microarray expression data using empirical Bayes methods. Biostatistics **8**, 118–127 (2007).

4. Leek, J. T., Johnson, W. E., Parker, H. S., Jaffe, A. E. & Storey, J. D. The sva package for removing batch effects and other unwanted variation in high-throughput experiments. Bioinformatics **28**, 882–883 (2012).
